# UPF1 localises to mitochondria and is linked to mitochondrial gene expression and function in Drosophila

**DOI:** 10.1101/2024.05.10.591322

**Authors:** Matthew T Wright, Hannah L Dixon, Parul Gopal, Emily M Price, Elizabeth Connolly, Alessandro Di Maio, Jonathon Barlow, Marco Catoni, Yun Fan, Anand K Singh, Saverio Brogna

**Affiliations:** School of Biosciences and Birmingham Centre for Genome Biology (BCGB), University of Birmingham, Edgbaston, Birmingham, B15 2TT, United Kingdom; Department of Biology, Indian Institute of Science Education and Research (IISER) Tirupati, Andhra Pradesh, India, 517619; Department of Biology, Faculty of Science, University of Copenhagen, Universitetsparken 15, Building. 3, 3^rd^Floor, 2100 Copenhagen O, Copenhagen, Denmark; Tech Hub Microscopy Facility, College of Medical and Dental Sciences, University of Birmingham, Edgbaston, Birmingham, B15 2TT, United Kingdom; Cellular Health and Metabolism Facility, School of Sport Exercise and Rehabilitation Sciences, University of Birmingham, Edgbaston, Birmingham, B15 2TT, United Kingdom

**Keywords:** Mitochondria, RNA helicases, NMD, UPF1, *Drosophila*

## Abstract

UPF1 is a conserved RNA helicase in eukaryotes typically defined by its role in nonsense-mediated mRNA decay (NMD). This study presents ChIP–seq evidence indicating an RNA-dependent interaction of UPF1 with mtDNA, consistent with an RNA-mediated association. Mitochondrial localisation was observed as discrete foci throughout the mitochondrial network, frequently in proximity to nucleoids, as confirmed microscopically by immunostaining and GFP tagging of UPF1 in different fly tissues and S2 cells. Depletion of UPF1, but not depletion of the other core NMD factors UPF2 or UPF3, in salivary glands results in smaller mitochondria and, unexpectedly, increased respiratory activity. This increase is observed specifically in salivary glands and correlates with elevated expression of nuclear genes involved in respiratory chain assembly and mitochondrial translation. Changes in mtDNA-encoded gene expression are also observed upon the depletion of UPF1. These findings indicate that UPF1 operates in mitochondria and, independently of canonical NMD, is linked to mitochondrial gene expression and may coordinate nuclear and mitochondrial gene expression in a tissue-specific manner.

## Introduction

UPF1 is an evolutionarily conserved RNA helicase in eukaryotes necessary for accurate gene expression. This helicase binds both RNA and single-stranded DNA *in vitro* and uses ATP hydrolysis to unwind secondary structures and displace associated proteins ^1–4^. UPF1 is predominantly cytoplasmic in most cells, where it associates with mRNA in a translation-dependent manner ^5–7^. Although best known for its role in nonsense-mediated mRNA decay (NMD), an mRNA surveillance pathway understood to recognise and degrade transcripts undergoing premature translation termination ^8–16^.UPF1 also shuttles rapidly between cellular compartments ^17–21^. In the nucleus, UPF1 associates with chromatin, and in mammalian cells it has been proposed to play a direct role in DNA metabolism ^22,23^. Studies in *Drosophila melanogaster* and *Schizosaccharomyces pombe* indicate that this association is primarily with nascent RNA and provide evidence that UPF1 may influence cotranscriptional steps of gene expression including release of the mRNA from the transcription site and its nuclear export ^20,24,25^. Misregulation of UPF1 has been linked to cancer and neurodegenerative disorders, including amyotrophic lateral sclerosis (ALS) and Alzheimer’s disease ^26–31^. However, it remains unclear whether these diseases as well as the multiple phenotypes attributed to UPF1 deficiencies are solely caused by cytoplasmic stabilisation of NMD targets or by yet unknown NMD-independent functions of UPF1.

The name UPF1 (*UP*-*F*rameshift-*1*) originates from its identification in a genetic screen for co-suppressors of an up-frameshift mutation in *Saccharomyces cerevisiae* ^32,33^. However, this gene was initially characterised as *NAM7* (*N*uclear *A*ccommodation of *M*itochondria *7*) in a screen for nuclear genes that, when overexpressed, restore respiration in strains carrying mutations in Group I or Group II introns of mitochondrial genes ^34,35^. Deletion of *UPF1*, *UPF2* and *UPF3*, which all equally inhibit NMD, leads to respiratory impairment in *S. cerevisiae*. However, only *UPF1* overexpression can restore mitochondrial splicing ^36^. Since UPF1 was not detected in *S. cerevisiae* mitochondria, these effects were interpreted to be a consequence of UPF1 overexpression, resulting in the misexpression of undefined nuclear-encoded proteins involved in mitochondrial splicing ^36^. However, the possibility that a small amount of UPF1 may localise to mitochondria and impact mtDNA gene expression could not be ruled out ^36^.

## Results

### UPF1 is associated with transcription sites and other regions of the mtDNA

Whilst analysing UPF1 ChIP-seq datasets generated during an earlier study ^20^, we made the unexpected observation that specific regions of the mtDNA are enriched and could therefore be directly or indirectly associated with UPF1. The enrichment is most apparent at the A-T-rich region of the mtDNA, with the first half showing greater enrichment than the second (Figure 1A). This region consists of two arrays of highly similar sequence repeats that can be classified into distinct sequence types, one located in the first half, and the other in the second, separated by non-repetitive sequences ^37^. The A-T-rich region corresponds to the D-loop in mammalian mtDNA, also known as the major non-coding region (NCR) ^38^. In *Drosophila*, this region is predicted to contain a unidirectional mtDNA replication origin in its centre, which advances in the direction of the rRNA gene locus ^39^. The A-T-rich region is flanked by two promoter/transcription start site (TSS) regions that correspond to two major divergent transcription units: one containing a cluster of seven protein-encoding genes (TU1), and the other comprising the small and large rRNAs (TU3) and the ND1 gene (Figure 1A) ^40^. The TSS regions of the other two divergent transcription units, located between the ND4 and ND6 genes, also appear to be enriched for UPF1 (Figure 1A; TU2 and TU4) ^40^. The enrichment profile is similar between the two biological replicates, despite differences in the overall levels of enrichment in the two repeats (Figure 1A, top red tracks). The enrichment peaks mostly map to the inter-repeat regions (Figure 1A, horizontal yellow arrows); particularly in the second half of the A-T region (a prominent inter-repeat peak is denoted by the black vertical arrow in Figure 1A). The regional specificity of the enrichment, together with the observation that mtDNA is not typically enriched in ChIP–seq datasets, suggests that this is unlikely to reflect background signal. For example, there is no evidence of enrichment of any mtDNA region in the Ser2 Pol II ChIP-seq dataset, which was generated in parallel, and similarly aligned, nor in other published datasets (Figure S1A). Likewise, enrichment of the AT-rich region does not appear to result from sequencing bias or alignment artefacts, sequencing of the input sample shows uniform coverage across most of the mtDNA, with somewhat lower coverage in the A-T region (Figure 1A, bottom purple graph).

**Figure 1.**
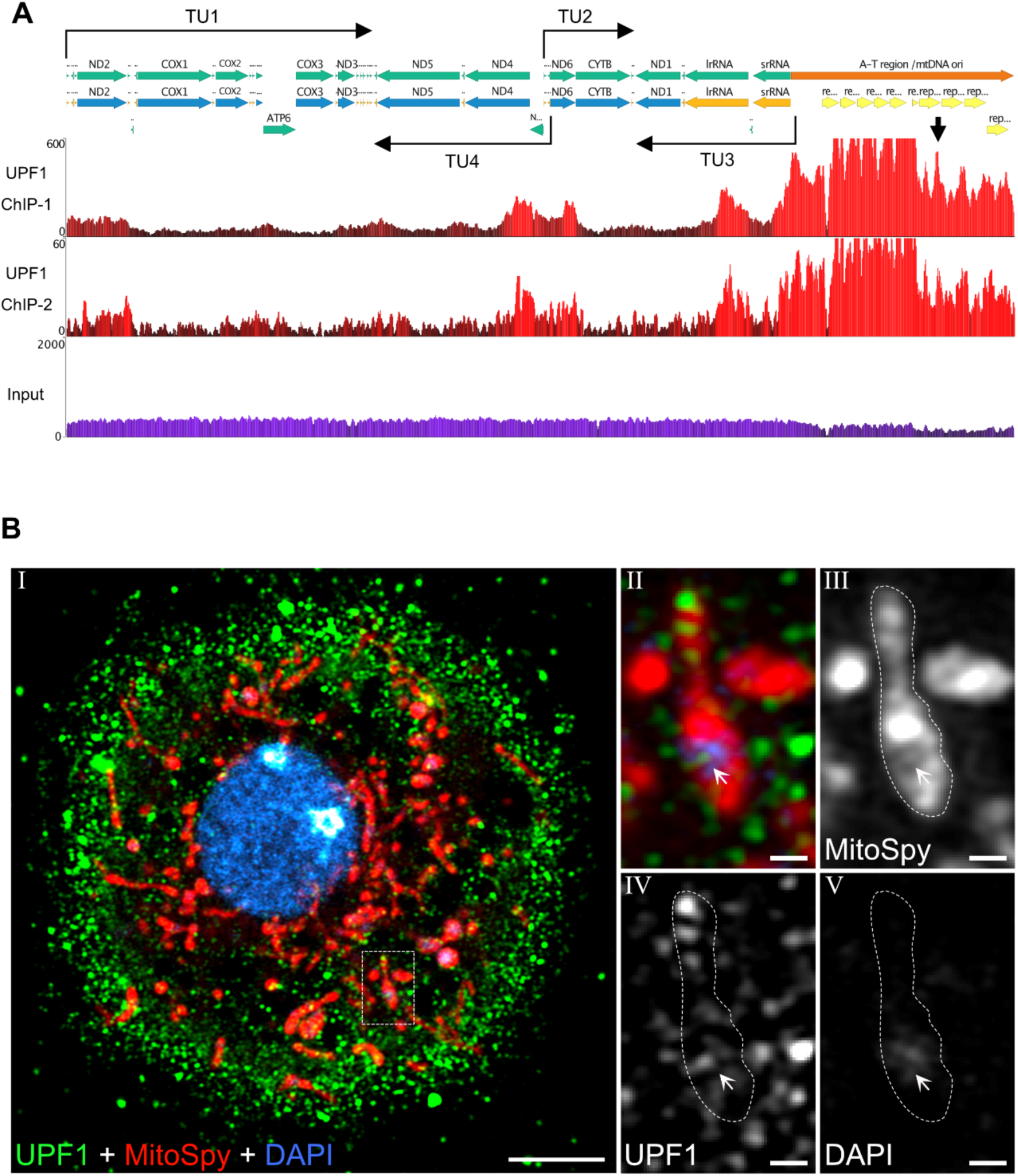
UPF1 mtDNA association and mitochondrial localisation in S2 cells. (**A**) The top schematic shows the linear map of the mtDNA and the location of the four transcription start sites and the corresponding transcription units (TUs), TU1 and TU2 on the top strand, TU3 and TU4 on the bottom strand. The protein-coding and rRNA genes are indicated in green, the A-T region / NCR is indicated in orange, blue indicates protein coding genes, orange indicates small and large rRNA, srRNA and lrRNA, respectively, yellow arrows indicate sequence repeats within NCR. The black arrow indicates a prominent peak of enrichment at one of the interrepeat regions. The bottom panel shows the genome browser enrichment profiles of two independent UPF1 ChIP-Seq experiments (red) and, as control, the profile of the input DNA (purple). The Y axes were not scaled equally, although the two UPF1 samples show different absolute levels of enrichment, the pattern is very similar. (**B**) Fluorescence immunolocalisation of endogenous UPF1 (green; using anti-UPF1 monoclonal antibody 7B12. A similar pattern is detected using 1C13, see Material and Methods) in S2 cells. The red signal corresponds to MitoSpy, a fluorescent dye that localises to the mitochondrial matrix (Materials and Methods). The blue signal corresponds to DAPI, which labels the nuclear DNA and the mtDNA. Panel I shows a merge image in the three channels of a whole cell: magnified views of the boxed region as shown in panels II – V, either as a merged image (panel II) or individual labelled channels in grey. The arrow points to a strong UPF1 signal dot adjacent to the DAPI stained region of the mitochondria. The images were taken using a Zeiss LSM900 confocal microscope equipped with a 63X objective lens (Materials and Methods). The scale bar corresponds to 10 µm.

The association with the rRNA gene loci and the ND4 region was validated by independent ChIP-qPCR experiments. These show an enrichment profile comparable to that observed in the ChIP-seq data (Figure S1B; the A-T region could not be tested as specific PCR primers could not be designed due to the high A/T content of the sequence). Additionally, the association with the ND4 and rRNA regions is RNase-sensitive (Figure S1C). Although the A-T region could not be assessed for RNase-sensitivity, mitochondrial transcription promoters are bidirectional, and this region is therefore likely to be highly transcribed from both the TU1 and TU3 promoters in the antisense directions. It is possible that the association of UPF1 with this region is also RNA dependent.

### Microscopic visualisation of UPF1 within mitochondria

UPF1 could be detected in mitochondria by immunostaining in S2 cells (Figure 1B). This revealed a punctate signal that is most intense in the cytoplasm, however some of the UPF1 foci are localised within the mitochondria, identified by staining cells before fixation with a fluorescently conjugated dye which accumulates in the matrix of functional mitochondria (Figure 1B) (MitoSpy; Materials and Methods). At this level of resolution it cannot be excluded that some UPF1 foci are associated with the mitochondrial surface; however, a subset colocalises with DAPI-stained regions (Figure 1B, panels II-V, magnified views of a mitochondrion; the arrow points to one of these foci), which likely correspond to nucleoids – clusters of multiple mtDNA molecules ^41^.

Purification of mitochondrial fractions from S2 cells, followed by western blotting also indicates that a portion of UPF1 co-purifies with cytochrome c oxidase subunit 4 (COX4), a component of the respiratory chain located in the inner mitochondrial membrane. In contrast, α-tubulin is barely detectable, indicating that these mitochondrial fractions contain minimal cytoplasmic contaminants (Figure S2).

Next, we examined whether UPF1 could be detected in salivary glands cells. A portion of UPF1 was detected within mitochondria by both immunostaining of endogenous UPF1 and imaging of a transgenic GFP-tagged UPF1. Immunostaining of endogenous UPF1 shows a profuse punctate signal in the cytoplasm and, to a lesser extent, within mitochondria - demarcated by COX4 signal in second instar larval glands (Figure 2A, the inset shows a magnified view of UPF1 signals within an intra-mitochondrial DAPI stained region). Mitochondrial localisation is also apparent in late third instar larval glands (Figure 2B, at this development stage, the cells are replete with large secretory vesicles).

**Figure 2.**
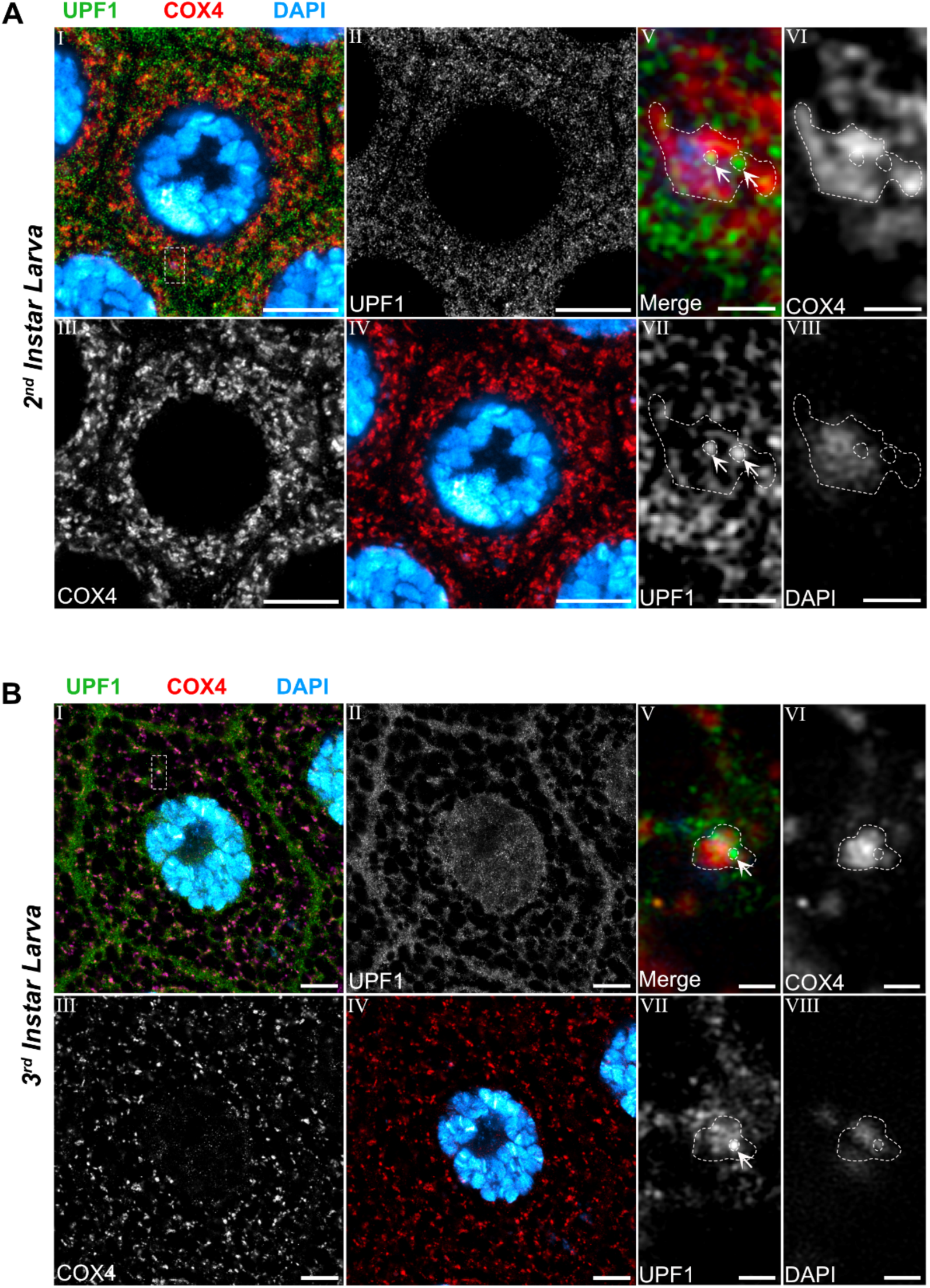
Endogenous UPF1 colocalises with COX4 in salivary glands. Fluorescence immunolocalisation of endogenous UPF1 (green) and COX4 (red) as the mitochondrial marker. DAPI (blue) represents DNA (both nuclear and mtDNA). (**A**) Shows a second instar larva salivary gland cell. (**B**) A third instar larvae cell. In both instances, panel I represents a merged image of all three channels, UPF1 (panel II), COX4 (panel III). Panel IV represents both COX4 and DAPI. The boxed region is shown magnified in panels V-VIII, as merged (V) and grey panels for COX4 (VI), UPF1 (VII) and DAPI (VIII). Arrows point to foci of UPF1 adjacent to DAPI-stained regions of the mitochondria. Scale bar represents 10 µm.

The transgenic GFP-UPF1, expressed using the UAS-GAL4 system with the salivary gland-specific driver *FkhGAL4*, (Materials and Methods), shows that a portion of the protein is also located within mitochondria (Figure S3). At high magnification, the glands display a dense GFP-UPF1 punctate pattern with foci of variable size and brightness that, although brighter overall in the cytoplasm, are also detectable within mitochondria (Figure S3, panels V-VIII show a strong GFP-UPF1 dot that colocalises with DAPI).

### Prominent UPF1 mitochondrial localisation in flight muscles

UPF1 mitochondrial localisation is prominent in adult flight muscles. These are characterised by distinctive rows of large, tubular, fused mitochondria, located between the myofibrils. Both immunostaining of endogenous UPF1 (Figure 3A) and imaging of transgenic GFP-UPF1 expressed in muscle (Figure 3B), using the muscle specific *MhcGAL4* driver (Materials and Methods), show that UPF1 appears to be relatively more abundant within mitochondria than in the cytoplasm. Regions of cytoplasm can be distinguished as spaces between myofibril bundles that are not occupied by mitochondria, such as the region indicated by the dashed yellow rectangle in Figure 3A, panels I-II. Bright UPF1 signal foci are often detected within mitochondrial regions that do not stain for COX4 (Figure 3A, panels IV-VI). These possibly correspond to nucleoids; however, this could not be verified by DAPI staining. Mitochondrial localisation is particularly apparent when GFP-UPF1 is expressed alongside the UAS-*mCherry-mito* transgene to demarcate the mitochondria (see Materials and Methods), where most of the protein appears predominantly localised within mitochondria, with only a small proportion present in the cytoplasm (Figure 3B, panels I-II).

**Figure 3.**
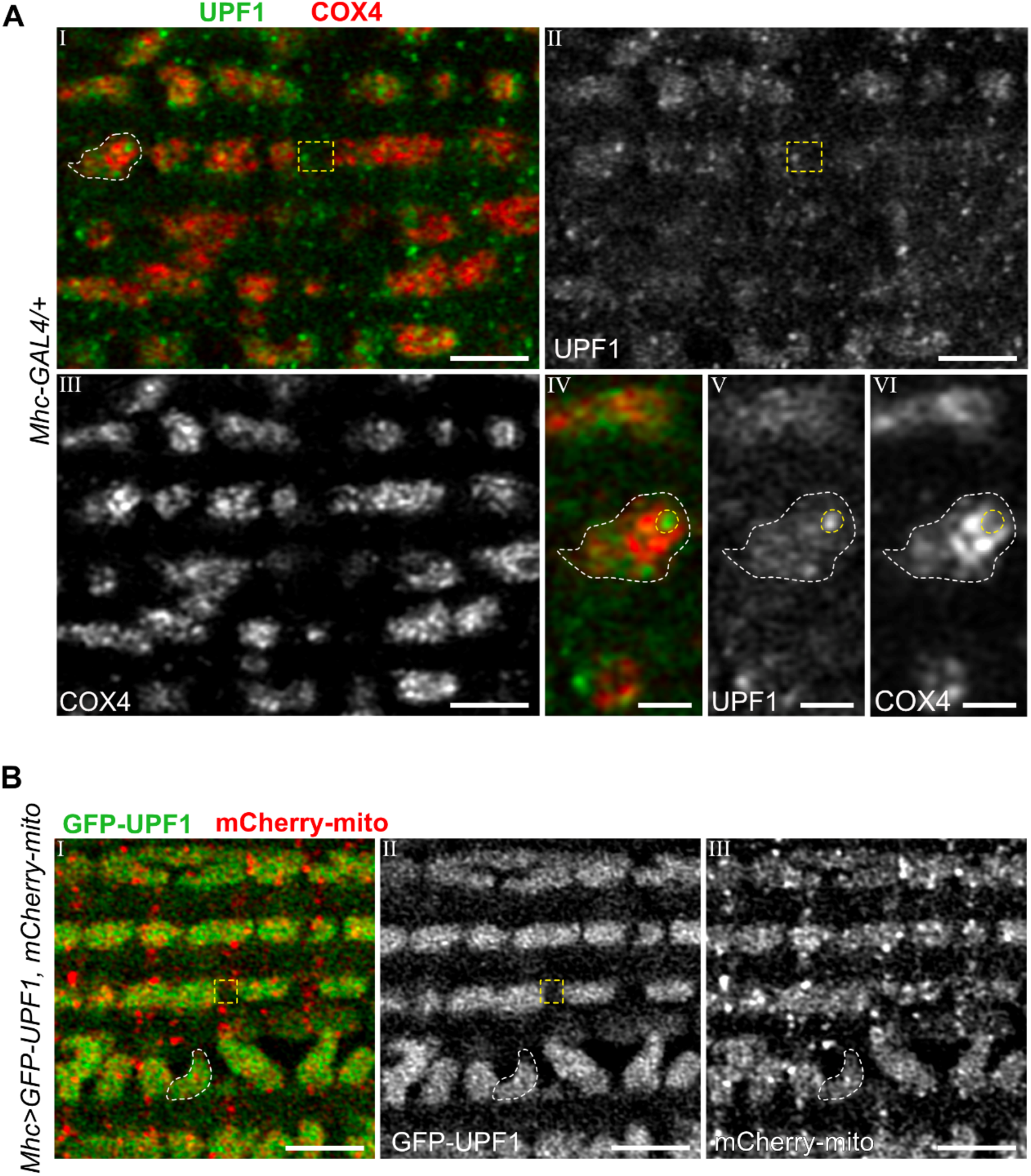
UPF1 displays prominent mitochondrial localisation in flight muscles. (**A**) Fluorescence immunolocalisation of endogenous UPF1 (green) and COX4 (red) as the mitochondrial marker in indirect flight muscles. Panel I shows a merged image of both channels, UPF1 (panel II) and COX4 (panel III), with an example cytoplasmic region indicated by the yellow dashed rectangle. Panels IV-VI represent a magnified view of the mitochondrial region outlined in panel I, as the merged image (panel IV), or as individual channels in grey, UPF1 (panel V) and COX4 (panel VI). A strong dot of UPF1 signal is circled in yellow in the magnified views; this is localising in a region lacking COX4 signal. (**B**) Fluorescence localisation of *MhcGAL4* driven expression of GFP-UPF1 (green) and mCherry-mito (red) in indirect flight muscles. Panel I represents a merged image of both channels, GFP-UPF1 (panel II) and mCherry-mito (panel III), a cytoplasmic region is indicated by the yellow dashed rectangle.

### UPF1 localisation in mitochondria is dynamic in spermatids

To further investigate UPF1 mitochondrial localisation, we examined *Drosophila* testes, as these contain cells with characteristic large mitochondrial structures. Spermatogenesis begins with a germline stem cell, which, after four cycles of mitosis, generates 16 primary spermatocytes, that produce 64 haploid spermatids following meiosis (Figure 4A). Both cell types display recognisable mitochondrial structures: the mitoball, a recently described transient bundle of mitochondria in premeiotic spermatocytes ^42^, and the classic nebenkern, a large spherical structure adjacent to the nucleus of round spermatids, formed by fusion of all mitochondria within these cells at this stage of development (the onion stage, Figure 4A) ^43^.

**Figure 4.**
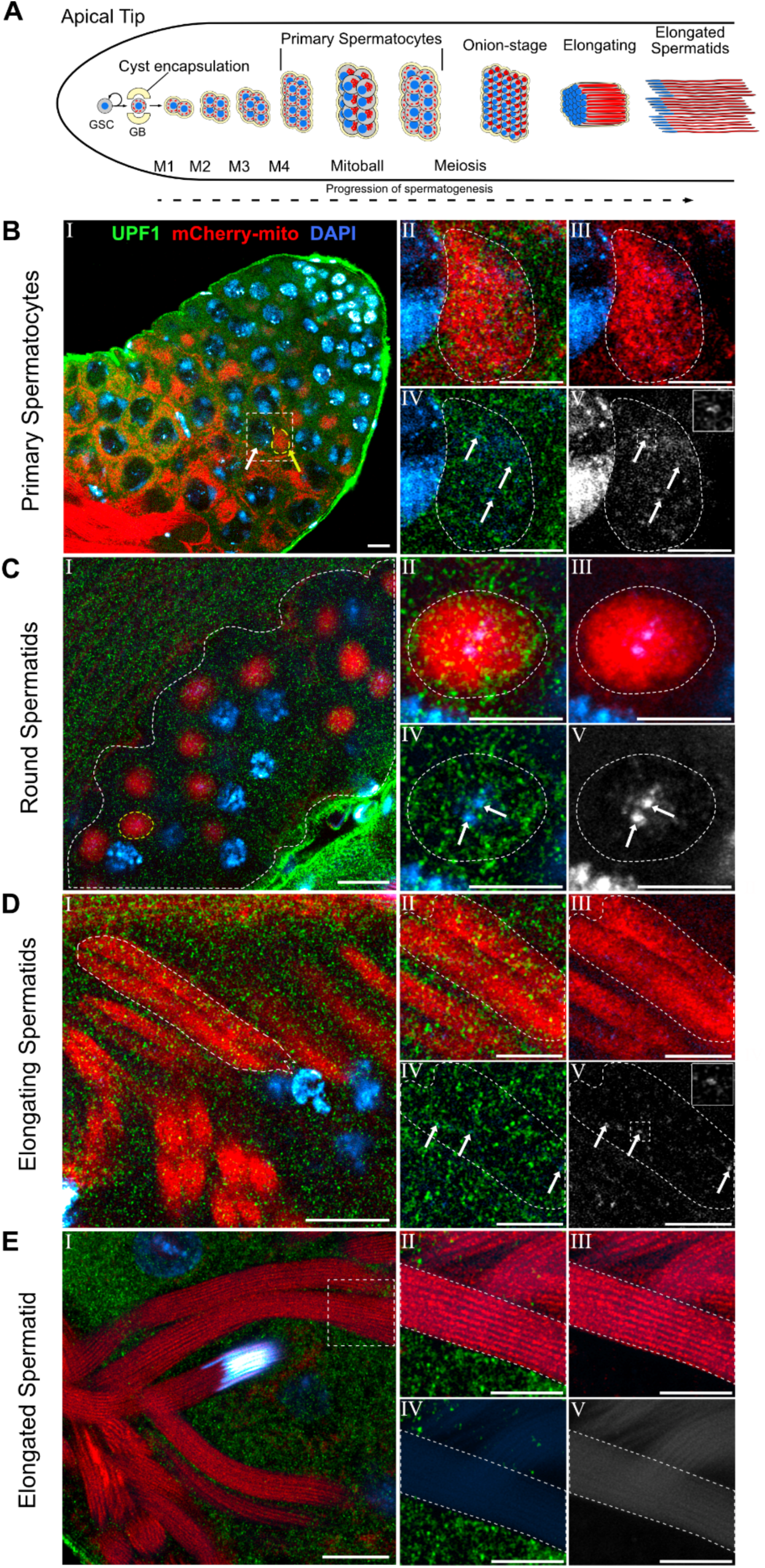
UPF1 mitochondrial localisation during spermatogenesis. (**A**) Schematic of *Drosophila* spermatogenesis. A germinal stem cell (GSC) at the apical tip of the testes proliferates into a differentiating gonialblast (GB), and a self-renewing GSC. The GB is enclosed by within a cyst made of two cyst cells and undergoes four rounds of mitosis (M1-M4) to produce 16 diploid primary spermatocytes. The mitochondria (red) cluster to form the Mitoball, a transient structure adjacent to the nucleus (blue). The mitoball disperses and the primary spermatocytes undergo meiosis to produce 64 haploid, round onion-stage spermatids, the mitochondria have fused to form the nebenkern adjacent to the nucleus. Each spermatid then undergoes elongation, with the unfurling of the nebenkern into the major and minor mitochondrial derivatives, and into the elongating tail. (**B-E**) Fluorescent immunolocalisation of endogenous UPF1 (green), imaging of ubiquitously expressed mCherry-mito (red) as the mitochondrial marker, and DAPI (blue) to stain nuclear and mitochondria DNA. (**B**) Primary spermatocytes. Panel I shows a merged image of the three channels, with a primary spermatocyte outlined with the white dotted box, the white arrow corresponds to the nucleus, and the yellow arrow marks the Mitoball. Panel II represents the merged magnified view of the outlined primary spermatocyte in panel I, the Mitoball is outlined in white. Panel III – IV represent mCherry-mito (panel III) and UPF1 (panel IV) together with DAPI. Panel V shows DAPI alone, with a magnified inset showing one nucleoid indicated by the topmost arrow (top right). White arrows in panels IV and V indicate UPF1 dots proximal to DAPI signal. (**C**) Shows a cyst with round-stage spermatids. Magnified views of the nebenkern outlined in yellow in panel I, are indicated in panels II-V. White arrows in panels IV and V point to two clusters of mtDNA/nucleoids in the centre nebenkern. (**D**) Localisation in elongating spermatids. Magnified view of a portion of the area (major and minor mitochondrial derivatives) outlined by the dashed line in panel I are shown in panels II – V. Arrows in panels IV and V show dots of UPF1 adjacent to DAPI, magnification inset of DAPI is seen in panel V (top right). Panel V shows DAPI alone. Arrows in panels IV and V indicate UPF1 dots proximal to residual DAPI foci. (**E**) Imaging of elongated spermatids, which show absence of UPF1. Panel I shows the outlined region which is magnified in panels II-V which show absence of UPF1 and DAPI foci in the spermatid tail, panels IV-V. Scale bars in B-E, panel I represent 10 µm, scale bars in B-E, panels II-IV represent 5 µm.

We first examined the subcellular localisation of endogenous UPF1 by immunostaining testes ubiquitously expressing mito-mCherry as a mitochondrial marker (*ubi-mito-mCherry*) ^44^. This shows punctate staining of UPF1, with a comparable level of signal in mitochondria and cytoplasm in most cell types (Figure 4B-D). This can be observed in both the mitoball (Figure 4B) and the nebenkern (Figure 4C). A similar pattern was seen in the axonemal mitochondria of elongating spermatids (Figure 4D). However, UPF1 signals are absent in the axonemal mitochondrial derivatives of fully elongated spermatids (Figure 4E, panels II and IV). These also lack DAPI foci in their tails (Figure 4E, panel V), consistent with the expected elimination of paternal mtDNA prior to full elongation ^45^.

We also examined the subcellular localisation of transgenic GFP–UPF1 alongside mito–mCherry using the cell-specific driver *bamGAL4:VP16* (*bamGAL4*), which is highly expressed in early spermatogonial cells ^46^. Similar to the endogenous protein, GFP–UPF1 also displayed a comparable mitochondria localisation pattern (Figure S4).

These observations indicate that UPF1 localises to mitochondria in both primary spermatocytes and developing spermatids, and that UPF1 signals are distributed throughout mitochondria, including nucleoid regions. The exception is fully elongated spermatids, where neither UPF1 nor nucleoid/mtDNA signals are present. The localisation of UPF1 to mitochondria therefore appears to correlate with the presence of mtDNA in spermatids.

### UPF1 depletion results in smaller mitochondria with increased respiration activity in salivary glands

Given the localisation of UPF1 in mitochondria, we next examined whether its depletion results in mitochondrial morphological changes in the cell types examined. Apparent changes were detected only in salivary glands. UPF1-knockdown (UPF1KD) salivary gland cells show smaller, more numerous mitochondria with less complex shapes in both L2 and L3 stages. This phenotype is visually apparent when using two independent RNAi lines (Figure 5A, panels II, III, V and VI) and is UPF1-specific, with no apparent phenotype observed in UPF2KD or UPF3KD salivary gland cells at either L2 or L3 stage (Figure 5B, panels I–IV).

**Figure 5.**
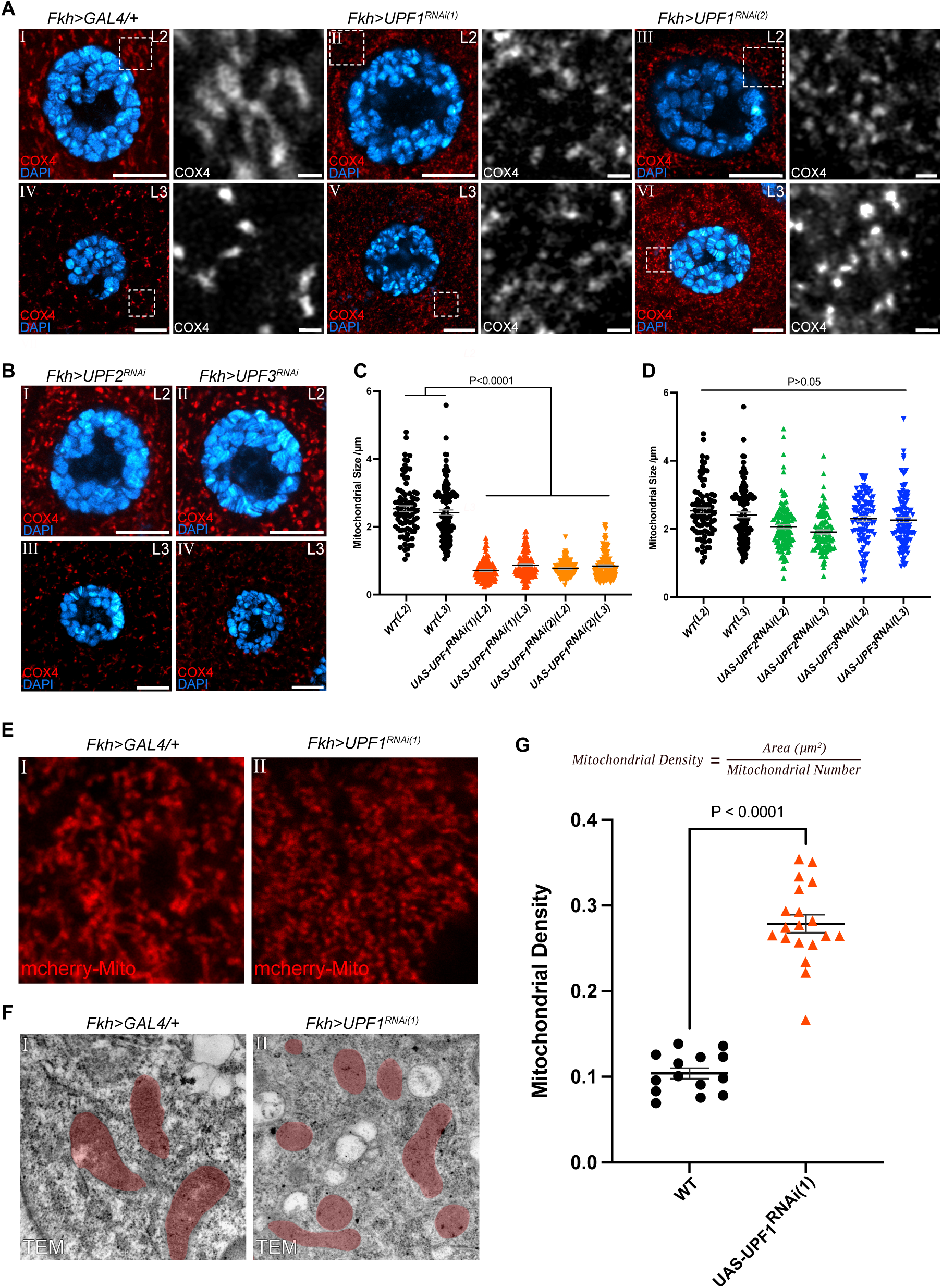
UPF1KD salivary glands have smaller but overall, more mitochondria. Imaging of mitochondria in WT, and UPF1, UPF2 and UPF3 depleted salivary gland cells in second (L2) and third instar (L3) larvae**. (A)** Immunofluorescent microscopy of salivary glands, COX4 (red) was used as the mitochondrial marker, DAPI (blue) stains DNA, both L2 and L3 salivary glands were examined. *FkhGAL4* was used as a control (WT; panel I and IV) and to induce depletion of UPF1 (two RNAi lines panels (II, V and V and VI for II & III, respectively)), UPF2 (panels VII and IX) and UPF3 (VIII and X). Magnified insets outlined in white (panels I-IV) show mitochondrial morphology to the right, in grey, apart from UPF2 and UPF3 RNAi (panels VII-X). (**B**) Immunofluorescent microscopy of mitochondria in UPF2 (panels I and III) and UPF3 (panels II and IV) depleted salivary gland cells in L2 and L3. (**C)** Measurement of mitochondrial size from mitochondria in WT and UPF1KD salivary gland cells in L2 and L3 larvae, (**D**) Measurement of mitochondrial size from mitochondria in WT, and UPF2 and UPF3 depleted salivary gland cells. Statistical significance was calculated using an ordinary one-way ANOVA with Šídák’s multiple comparisons test. The means were compared within L2 and L3 groups, but not between. Error bars represent the SEM. (**E**) Mitochondrial morphology imaging using mCherry-mito (red) as the mitochondria in WT (panel I) and UPF1KD (panel II) salivary gland cells. (F) TEM microscopy of WT (panel I) and UPF1KD (panel II) salivary gland cells. Red regions depict mitochondria. (**G**) TEM quantification of mitochondrial density (area (µm^2^) ÷ mitochondrial number) in WT and UPF1 depleted salivary gland cells. Significance was calculated using a two-tailed unpaired t-test, error bars represent the SEM. RNAi efficiency was estimated by RT-qPCR using gene-specific primers. RNA was reduced to 30-40 % of control for all RNAi lines used.

Mitochondrial size variation was quantified by measuring mitochondrial length. This analysis indicates that mitochondria are on average 2.53 µm in wild type, with a significant decrease to 0.73 µm in UPF1KD cells, and minimal change in size between the L2 and L3 developmental stages (Figure 5C). No significant change in mitochondrial size was detected in UPF2KD or UPF3KD salivary glands (Figure 5D).

This phenotype was further validated by expression of mito–mCherry and visualisation of mitochondria in wild-type and UPF1KD salivary glands (Figure 5E). Additionally, transmission electron microscopy (TEM) imaging of wild-type and UPF1KD salivary glands confirmed that reduced UPF1 results in smaller (Figure 5F) and significantly more numerous mitochondria and increased mitochondrial density (Figure 5G).

Next, we assessed mitochondrial function following RNAi mediated depletion in salivary glands by evaluating mitochondrial respiration using high-resolution respirometry (see Materials and Methods). This approach allows assessment of the function of individual electron transfer complexes and determination of whether respiratory activity is coupled to efficient ATP production.

Respiratory complexes I, II and IV were examined in homogenised salivary glands from wild type and UPF1KD. The activity of complexes II and IV, but not complex I, was significantly increased in UPF1KD compared to wild type (Figure 6A), with the greatest increase observed for complex IV. These changes were accompanied by an increase in succinate-dependent OXPHOS respiration (Figure 6B). When analysed under substrate-specific conditions that probe electron entry into the electron transport chain, only succinate-driven (complex II–dependent) respiration was increased, whereas NADH-linked (complex I–dependent) respiration remained unchanged, indicating enhanced electron input via the succinate pathway. Complex II–dependent respiration remained efficiently coupled to proton flux across the inner mitochondrial membrane, as indicated by increased respiratory control ratios and coupling efficiencies (Figure 6C–D). In contrast, direct measurement of complex IV under saturating conditions, independent of upstream electron supply, revealed a marked increase in oxygen consumption compared to wild type (Figure 6E), consistent with increased capacity at the terminal step of the electron transport chain. Together, these data indicate that the smaller mitochondria in UPF1KD are functionally active and exhibit increased respiratory activity compared to wild type. The precise mechanism underlying the increased respiratory activity was not investigated.

**Figure 6.**
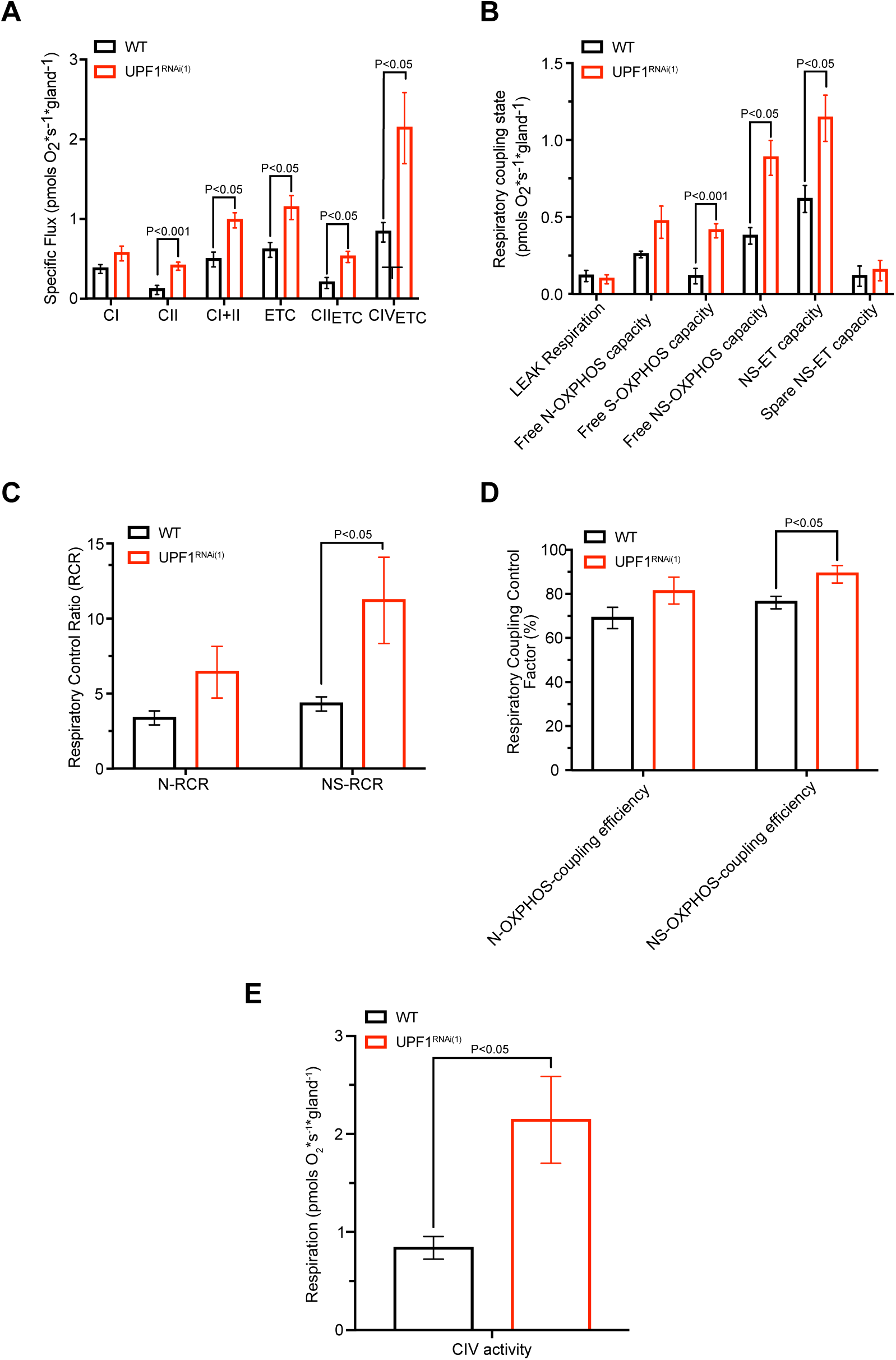
Mitochondrial respiration is elevated in UPF1KD salivary glands. (**A**) Specific oxygen flux (pmol O₂ s⁻¹ gland⁻¹) in control (green) and UPF1KD (red) salivary glands under substrate-defined respiratory conditions: complex I-linked (CI), complex II-linked (CII), combined complex I+II (CI+II), and uncoupled electron transport system capacity (ETS; CI+II). CII-driven ETS capacity (CIIETS) and complex IV capacity (CIVETS) are also shown. (**B**) Respiratory states (pmol O₂ s⁻¹ gland⁻¹) under LEAK and OXPHOS conditions. (**C**) Respiratory control ratios (RCR) for complex I (N-RCR) and combined complex I+II (NS-RCR) substrates. (**D**) Coupling efficiency of oxidative phosphorylation for complex I (N-OXPHOS) and complex I+II (NS-OXPHOS) substrates. (**E**) Complex IV respiratory capacity (pmol O₂ s⁻¹ gland⁻¹), measured in the presence of ascorbate and TMPD. Data are presented as mean ± SEM from three independent biological replicates. Statistical significance was assessed using unpaired t-tests (E) or multiple unpaired t-tests (A-D).

### Upregulation of respiration and mitochondrial translation genes in UPF1KD

Next, we examined whether there are transcriptomic changes that could explain the observed mitochondria phenotypes in salivary glands. RNA-seq was performed on UPF1KD, UPF2KD, and UPF3KD salivary glands alongside the wild-type as control (Figure 7A). In UPF1KD samples, upregulated genes are enriched within mitochondrial organisation, mitochondrial gene expression, mitochondrial translation, and respiratory chain complex assembly GO terms (Figure 7B). Notably, several of the enriched categories specifically relate to mitochondrial protein synthesis and the assembly of oxidative phosphorylation (OXPHOS) complexes, suggesting a coordinated increase in mitochondrial translational capacity and respiratory machinery. This upregulation provides a possible explanation for the increased mitochondrial respiration observed in UPF1KD salivary glands. Downregulated genes are enriched for amino acid catabolic processes (Figure 7B, right panel), indicating altered expression of pathways involved in amino acid metabolism. Both UPF2KD and UPF3KD salivary glands did not demonstrate functional enrichment of genes within mitochondria-associated GO terms (Figure S5A and B, respectively).

**Figure 7.**
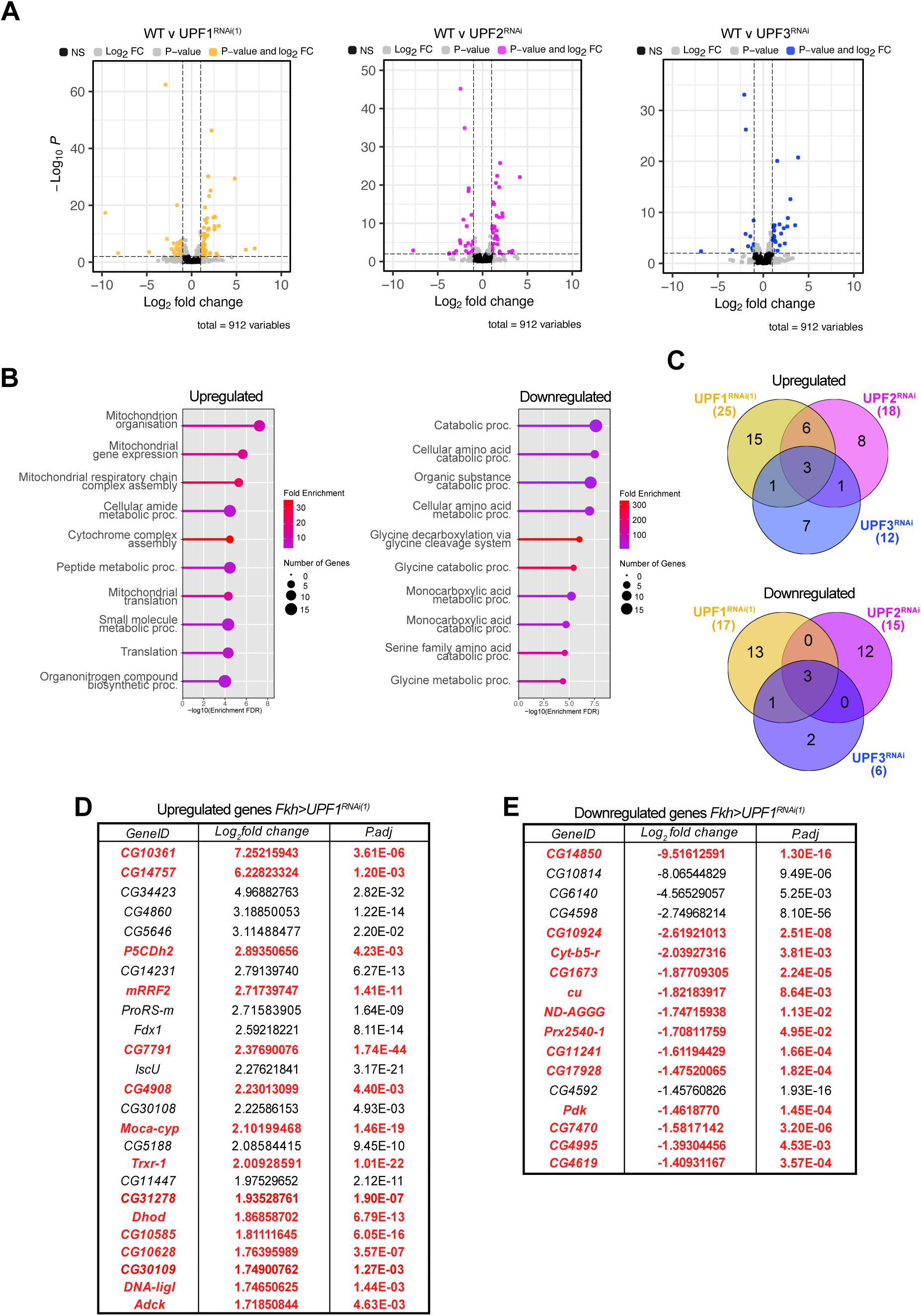
UPF1KD in salivary glands increases expression of genes involved in mitochondrial processes. (**A**) DESeq2 differential expression analysis of mitochondrial genes (n = 912; GO:0005739), in UPF1KD (red), UPF2KD (orange) and UPF3KD (green) vs WT salivary gland RNA-seq samples. Coloured points correspond to genes which are significantly differentially expressed (-1<Log_2_ fold change>1, P<0.05), those in grey are either significant but not highly misregulated (P<0.05; 1> Log_2_ fold change <-1), or misregulated but not significant (-1<Log_2_ fold change>1; P>0.05) those in black are not significant nor misregulated (P>0.05; 1> Log_2_ fold change <-1). (**B**) GO analysis of UPF1KD salivary glands was performed on significantly misregulated genes with the enriched upregulated (left panel) and downregulated (right panel) biological processes shown. The node size corresponds to the number of genes associated with the GO term, colour corresponds to the fold enrichment, from blue (lowest) to red (highest), only pathways with a -log10(enrichment FDR) <0.05 are shown. Pathways are ordered by -log10(enrichment FDR). (**C**) Venn diagrams showing the number of shared and unique upregulated (top panel) and downregulated (bottom panel) mitochondria-associated genes for all three genotypes. (**D-E**) List of the top upregulated (D) and downregulated (E) genes (Log_2_ fold change >1; p-adjusted (P.adj) <0.05). Genes highlighted in red are exclusively misregulated in UPF1KD.

Specifically, 15 nuclear genes encoding mitochondrial proteins are selectively upregulated in UPF1KD glands, highlighted in red in Figure 7D. Of the upregulated genes, CG14757 (Succinate dehydrogenase assembly factor 2-A, mitochondrial; Sdhaf2) is involved in the assembly of succinate dehydrogenase (respiratory complex II) ^47,48^. This may help to explain the observed increase in succinate-linked respiration. In addition, mRRF2 encodes a mitochondrial ribosome recycling factor involved in mitochondrial translation ^49^, and CG31278 is predicted to encode a mitochondrial peptide deformylase required for processing of nascent mitochondrial translation products (FlyBase). CG10628 is predicted to be involved in ribosome maturation (FlyBase). Although not UPF1-specific, ProRS-m, a mitochondrial prolyl-tRNA synthetase, is also upregulated (log₂ fold change: 2.72 in UPF1KD, 1.69 in UPF2KD and 1.02 in UPF3KD glands; P.adj: 1.64 × 10⁻⁹, 4.21 × 10⁻⁵ and 4.04 × 10⁻², respectively). In contrast, 13 genes are specifically downregulated in UPF1KD glands (Figure 7C and 7E). Among these, Pdk (pyruvate dehydrogenase kinase) was identified. PDK phosphorylates and inhibits the pyruvate dehydrogenase complex, thereby limiting conversion of pyruvate to acetyl-CoA and entry of carbon into the tricarboxylic acid cycle ^50^. Downregulation of Pdk would therefore be expected to increase acetyl-CoA production and promote respiratory flux.

Notably, the most downregulated gene (9.52 log₂ fold change) in the UPF1KD, CG14850, was recently characterised as Jig, a protein that shuttles between the mitochondria and the nucleus and, by binding the transcription factor CREB (CrebA), regulates the transcription of nuclear mitochondrial-associated genes in salivary glands ^51^. Depletion of Jig also results in smaller but active mitochondria. The downregulation of this retrograde signalling component suggests that UPF1KD alters mitochondria–nuclear communication, potentially contributing to the coordinated upregulation of mitochondrial translation and respiratory capacity. Both Jig downregulation and the smaller mitochondria phenotype cannot be explained by NMD inhibition, as these are not observed in UPF2KD and UPF3KD glands.

### Evidence of reduced expression of mtDNA genes in UPF1KD cells

Given the evidence for UPF1 association with mitochondrial transcription sites, we next examined whether there were transcriptomic changes in expression of the mitochondrial DNA in UPF1 depleted cells. The analysis revealed that six out of the 15 mtDNA genes are modestly, but statistically significantly, downregulated in UPF1KD S2 cells (Figure 8A and D). Although the magnitude of downregulation is small (between –0.2 and –0.3 log₂ fold change), the adjusted P-values indicate significant misregulation (P.adj ≤ 0.005). The strongest downregulated gene is ND4 (LFC: –0.29/–0.30; P.adj: 4.89E–03), which is the gene that showed the greatest UPF1-CHIP association in S2 cells

**Figure 8.**
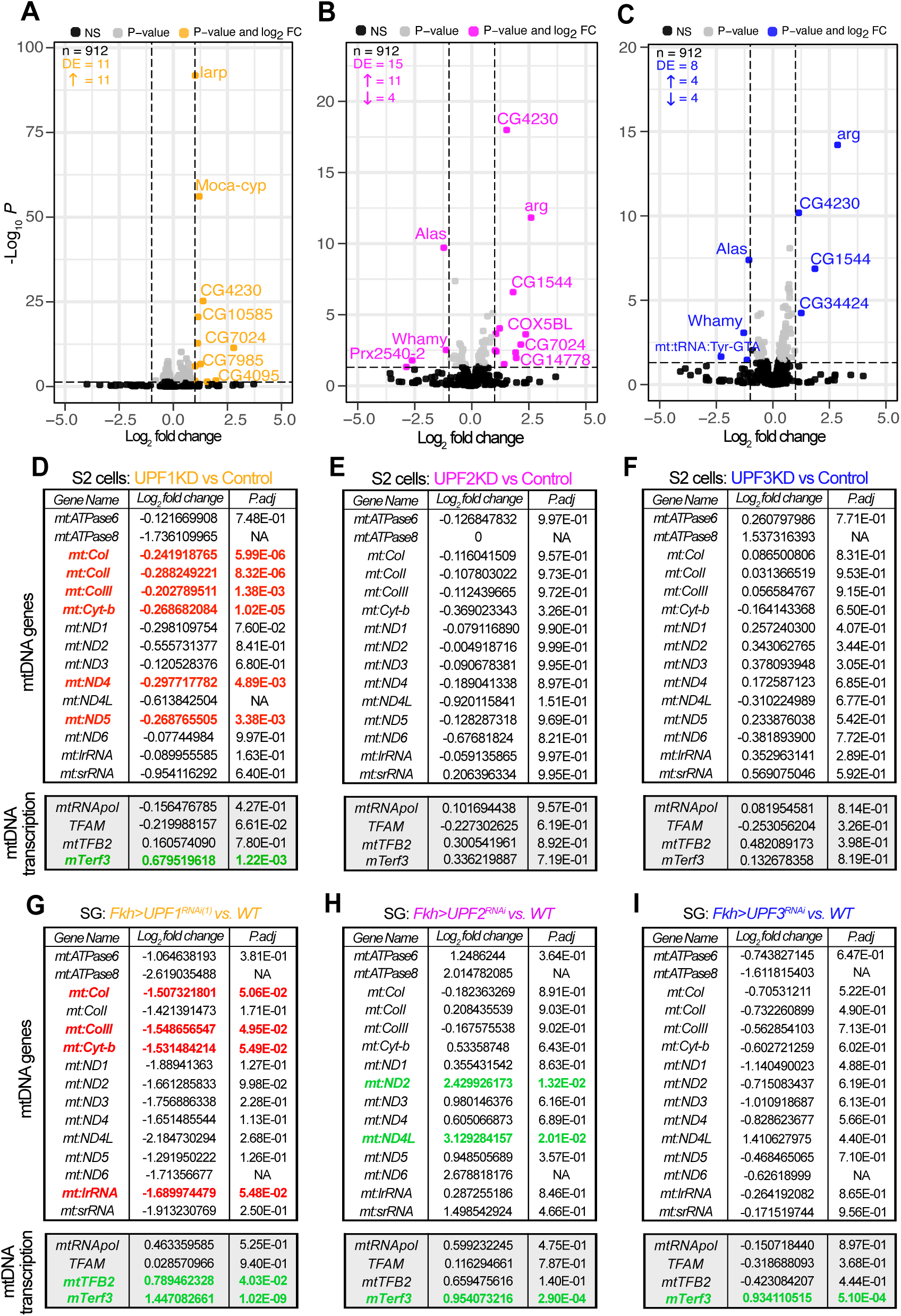
Indications of reduced mtDNA gene expression in UPF1KD S2 cells and salivary glands. (**A**) DESeq2 differential expression analysis of mtDNA genes n = 15, values were calculated based on the comparison against 17610 expressed genes, in S2 cells. (**A**) UPF1KD, (**B**) UPF2KD and (**C**) UPF3KD vs. control RNA-seq samples. Coloured points represent significantly differentially expressed mitochondrial genes (-1< LFC >1, P<0.05) with coloured dots: in the UPF1KD (orange; 11 upregulated and 0 downregulated), in the UPF2KD (magenta, 11 upregulated and 4 downregulated) and UPF3KD (blue, 4 upregulated and 4 downregulated) grey indicates significant, but not highly misregulated (1> LFC >-1, P<0.05) and those in black are non-significant (P>0.05). The gene ID, LFC and P.adj values are shown for the DESeq2 analysis of UPF1KD (**D**) UPF2KD (**E**) and UPF3KD (**F**) in S2 cells. The same is shown for the DESeq2 differential expression analysis of mtDNA genes n = 15 compared against 17284 genes in UPF1KD (**G**), UPF2KD (**H**) and UPF3KD (**I**) vs control RNA-seq salivary gland (SG) samples. Downregulated and upregulated genes associated with P.adj value of ≤0.05 are highlighted in red and green, respectively. For each panel (D-I), the list on the top shows mtDNA genes, while that at the bottom represents the 4 nuclear-encoded genes involved in mtDNA transcription (*mtRNApol*, *TFAM*, *mtTFB2* and *mTerf3*).

No significant changes in mtDNA gene expression were detected in UPF2KD (Figure 8B and E) or UPF3KD cells (Figure 8C and 8F). Additionally, there is no evidence of downregulation of genes involved in mitochondrial transcription in UPF1KD cells; instead, one of them, mTerf3 is upregulated (Figure 8, bottom panel).

Analysis of two independent RNA-seq datasets from the larval salivary glands indicates possible downregulation of several mtDNA genes (–1.5 < LFC), with four genes showing P.adj values around 0.05 (Figure 8G). Although these changes are just close to the statistical significance threshold, RNA-seq analysis of UPF2KD and UPF3KD salivary glands shows no evidence of mtDNA transcription downregulation (Figure 8H and 8I). Instead, UPF2KD salivary glands exhibit upregulation of two mtDNA genes (Figure 8H). The downregulation is unlikely to result from misregulation of nuclear genes required for mtDNA transcription, as none of these genes are downregulated. Rather, mtTFB2 and mTerf3 are upregulated in UPF1KD (Figure 8G), and mTerf3 is also upregulated in UPF2KD and UPF3KD salivary glands (Figure 8H and I, respectively).

## Discussion

We provide evidence that the RNA helicase UPF1 enters mitochondria and associates with mtDNA. This conclusion is supported by ChIP-seq data from Drosophila S2 cells demonstrating association of UPF1 with mtDNA transcription sites and possibly the major non-coding region. At gene loci, the association is most apparent at promoter-proximal regions and likely reflects interaction with nascent RNA, as indicated by RNase-sensitive ChIP–qPCR experiments. UPF1 was also detected within mitochondria by immunostaining and by expression of a transgenic GFP-tagged UPF1 in different Drosophila cell types. UPF1 signals are distributed throughout mitochondria, with some frequently colocalising with mtDNA. In most cells examined, UPF1 is less abundant in mitochondria than in the cytoplasm; however, in flight muscles and spermatids, mitochondrial levels are either greater than or comparable to those in the cytoplasm.

This localisation was unexpected, as the UPF1 sequence lacks an N-terminal mitochondrial targeting signal (MTS) and is not predicted to localise to mitochondria by computational methods ^52–54^. However, several core mitochondrial proteins similarly lack a classical MTS, and their localisation cannot be reliably inferred from primary sequence alone ^55^

Depletion of UPF1 resulted in reduced mitochondrial size and altered respiratory function in salivary glands. UPF1 knockdown was associated with increased respiratory activity and improved coupling efficiency, consistent with enhanced activity at both the entry (complex II) and terminal (complex IV) steps of the electron transport chain. The reduction in mitochondrial size may reflect a primary mitochondrial perturbation caused by UPF1 depletion, whereas the increased respiratory activity likely represents a compensatory response. Similarly, recent work in a murine pancreatic cancer model has linked UPF1 deficiency to increased complex I function and reactive oxygen species production. This resulted in immunosuppressive microenvironment and enhanced tumour growth ^31^. However, in our system, complex I activity is not affected.

Given the evidence for RNA-dependent association with mtDNA, UPF1 may directly modulate mitochondrial gene expression and play a role in mitochondrial transcription and transcription-coupled processes. Analogous to its role in the cytoplasm, UPF1 may influence mitochondrial mRNA stability and, given the upregulation of proteins involved in mitochondrial translation in UPF1KD, also affect mitochondrial translation. Perturbation of mitochondrial gene expression could be the primary trigger leading to reduced mitochondrial size which then trigger a compensatory mechanism involving upregulation of nuclear genes associated with respiration to maintain the respiratory output of the glands.

An alternative explanation is that the mitochondrial phenotype arises indirectly from altered expression of nuclear genes involved in mitochondrial function or feedback mechanisms that maintain coordination between nuclear and mitochondrial gene expression. In this scenario, UPF1 depletion would primarily affect nuclear gene expression, with secondary consequences for mitochondrial morphology and respiration. Notably, the reduction in mitochondrial size is observed in salivary glands but not in other tissues examined, suggesting that this effect is tissue-specific and that the underlying mechanism is likewise tissue-specific. This could be linked to Jig being expressed only in the salivary glands among the tissues examined, which, as mentioned above, has been proposed to play a role in mitochondrial–nuclear crosstalk and is drastically downregulated upon UPF1 depletion. ^51^.

Neither morphological nor transcriptional changes that could be linked to changes in respiration were observed in UPF2KD or UPF3KD glands, indicating that the phenotype is unlikely to result from general inhibition of NMD and instead reflects a specific function of UPF1.

Together, these findings link UPF1 to mitochondrial function in Drosophila and suggest that it may participate in the coordination of mitochondrial, nuclear and cytoplasmic gene expression.

## Materials and Methods

### Cell Culture, RNA interference and immunostaining

S2 cells (CVCL_Z232) were cultured in Schneider’s insect cell culture media (Catalogue number: ESF291, Generon) supplemented with 10% Fetal Bovine Serum (FBS), and 1% Penicillin-Streptomycin-Glutamine mix (P/S/G, Invitrogen) at 27°C. DNA for dsRNA generation were amplified by PCR using primers for the desired sequence in addition to the T7 promoter sequence at their 5’ end (bold) (5’-**TTAATACGACTCACTATAG**GGGAGA-3’). The PCR product was purified using the Monarch PCR and DNA cleanup kit (T1030S, NEB) and dsRNA was synthesised using the T7 RiboMAX express system (P1700, Promega). In a 6-well plate, 10^6^ cells were seeded in each well, incubated overnight, and then treated with 15 µg of dsRNA per well, in serum-free media. The cells were incubated for 1 hour at room temperature before being supplemented with 2 mL of complete media. The cells were incubated for three days to knockdown the UPF1 mRNA, before being harvested. For immunostaining, S2 cells (1 mL corresponding to about 10^6^ cells) were seeded onto Concanavalin-A (500 µg/mL) treated coverslips, in a 6-well plate for 30 minutes at 27°C to allow attachment. Excess media was eluted off and replaced with serum-free media containing typically 250 nM MitoSpy^TM^ Red CMXRos (424801, BioLegend) (in some experiments MitoTracker was used instead, as indicated in the figure legends; MitoTracker™ Red CMXRos, M7512, ThermoFisher Scientific). The cells were incubated with MitoSpy for 30 minutes at 27 °C, protected from light. Cells were then washed twice in serum-free media, fixed for 15 minutes in 4% paraformaldehyde at room temperature, washed twice in 1X PBS (13 mM NaCl, 0.7 mM Na_2_HPO_4_, pH 7.4), permeabilised for 15 minutes at room temperature in 0.1% PBST (1X PBS, 0.1% Triton X-100 T8787, Sigma-Aldrich), and blocked for 1 hour at room temperature in blocking solution (5% Bovine Serum Albumin (BSA), 0.05% sodium azide, and 1X PBS). They were then incubated in primary antibody for 16 hours at 4°C, washed and incubated in the appropriate, fluorophore-conjugated secondary antibody for 2 hours at room temperature. After incubation with antibodies, samples were washed in 1X PBS and incubated with DAPI (4-6-diamidino-2-phenylindole, Sigma-Aldrich, 1 µg/mL) for 10 minutes at room temperature. Samples were briefly washed in 1X PBS and mounted in VectaShield® antifade mounting medium (Vector Laboratories).

### *Drosophila* stocks and husbandry

Flies were reared on standard cornmeal fly food media at 25°C. The *y w^1118^* strain was used as wild type (DGGR_108736). The following lines were obtained from Bloomington *Drosophila* stock centre: *UAS-GFP-Upf1* (24623), *UAS-Upf1^HMC05537^* (referred to as UPF1^RNAi(1)^; 43144), *UAS-Upf1^GL01485^* (referred to as UPF1^RNAi(2)^; 64519), *UAS-mCherry-mito* (66533), *UAS-Upf2^JF01560^* (31095) and *UAS-Upf3^HMJ22158^* (58181). The following *Gal4* drivers *forkhead* (*Fkh)*^56^*, myosin heavy chain* (*Mhc*)^57^, *bag-of-marbles* (*bam*)^46^, drove expression of UAS constructs within the salivary glands, muscles, and late spermatogonia/early spermatocytes, respectively. The *ubi-mCherry-mito* is a transgene controlled by the Ubiquitin promoter ^44^. Combination stocks of second and third chromosomal transgenes were generated through standard combination crosses with a double balancer line (*w; IF/CyO-Act>GFP; MKRS/Tm6, Tb)*.

### Antibodies

Antibodies used for immunostaining: rabbit anti-COX4 (COX4I1, NB110-39115, Bio-Techne, diluted 1:100). The mouse monoclonal anti-UPF1 antibodies used was typically 7B12 or in some instances 1C13 (diluted 1:100). The same anti-UPF1 antibodies were used in ChIP-seq as described previously ^20^.

### S2 Cell fractionation and western blotting

To isolate the nuclear and cytoplasmic fractions, 2×10^6^ S2 cells were harvested and washed twice with ice-cold 1X PBS. Cells were resuspended in 1 mL ice-cold buffer AT (15 mM HEPES pH 7.6, 10 mM KCl, 5 mM MgOAc, 3 mM CaCl_2_, 300 mM Sucrose, 0.1% Triton X-100, 1 mM DTT, 1X PhosSTOP (4906845001, Roche), 1X cOmplete, Mini, EDTA-free Protease Inhibitor Cocktail (04693159001, Roche) and 1 U/mL Ribolock RNase Inhibitor) and incubated on ice for 10 min as a hypotonic treatment. Cells were lysed using a 2 mL Dounce homogenizer by applying 30-40 strokes with the tight pestle at 4°C. Lysates were divided into two aliquots of 500 µL and each aliquot was layered over 1 mL cushion of buffer B (15 mM HEPES-KOH at pH 7.6, 10 mM KCl, 5 mM MgOAc, 3 mM CaCl_2_, 1 M sucrose, 1 mM DTT, 1X cOmplete, Mini, EDTA-free Protease Inhibitor Cocktail). Following the centrifugation at 8000 RPM for 15 min at 4°C, the top layer (300 µL) was collected as cytoplasmic fraction. The pellet was resuspended in 500 µL AT buffer and layered over again on buffer B. Following the centrifugation, the pellet was collected as nuclear fraction. Proteins fractions were analysed by standard western blotting.

### Adult and larval tissue immunostaining

Internal organs of third instar larvae were dissected in 1X PBS, fixed for 20 minutes at room temperature in 4% paraformaldehyde (methanol-free; 43368.9L, VMR international), washed twice in 1X PBS, and permeabilised in 0.1% PBST (1X PBS, 0.1% Triton X-100). Samples were incubated in blocking solution for 2 hours at room temperature, antibody incubation, and sample mounting remain the same as mentioned above.

Testes of 2–5-day old adult males were dissected in 1X PBS. The only notable changes compared to larval tissue was a fixation time of 30 minutes at room temperature on a rocker, and one-hour permeabilization following fixation in 0.3% PBST (1X PBS, 0.3% Triton X-100) at room temperature. All other steps remained the same as mentioned prior. Testes squash preparations were prepared as described previously ^58^

For female muscle dissection, the previously established protocol was followed ^59^ and the samples were mounted in the same mounting media as mentioned for the prior cell and tissue samples.

### Confocal imaging and image analysis

The images of developing spermatids and nebenkern structure were captured under the phase contrast microscopic setting using x63, 1.4-NA oil immersion objective of Nikon Eclipse 80i fluorescence microscope. Confocal z-stack images of mounted samples were acquired using the Zeiss LSM900 with AiryScan2 module, Zeiss LSM710, Leica TCS SP8 STED, and Leica TCS SP2-AOBS confocal microscopes using Plan-Apo 10X, 25X, 63X, 1.4-NA objectives. All objectives, except for 10X, were oil immersion. Images were processed and quantified using FIJI ^60^.

Quantification of Total Corrected Fluorescence (TCF) was calculated using FIJI ^60^. A mean intensity projection of multi-z-stack image was generated, a region of interest was defined, and then the measure function was used. This was repeated for a background region to identify the mean of background fluorescence (MBF). The output was inserted into the following formula: TCF = Integrated Density – (Area * MBF).

### ChIP-Seq

Chromatin immunoprecipitation (ChIP) experiments were performed as previously described, ^20^. Briefly, S2 cells (2 x 10^7^) were washed and fixed in 1% formaldehyde (EM grade, Polyscience) for 10 minutes at room temperature. Cross-linking was stopped by addition of 125 mM Glycine for 5 minutes at room temperature. Cells were briefly centrifuged at 4°C, and rinsed in 1X PBS containing 1X cOmplete, Mini, EDTA-free Protease Inhibitor Cocktail. The pellet is resuspended in lysis buffer (5 mM PIPES pH 8.0, 85 mM KCl, 0.5% NP-40) supplemented within the same protease inhibitor cocktail and 1X PhosSTOP. Cells were centrifuged again, and rediluted in nuclear lysis buffer (50 mM Tris pH 8.0, 10 mM EDTA, 1.0% SDS) with protease inhibitor cocktail and PhosSTOP. The cell suspension is diluted with IP dilution buffer (16.7 mM Tris pH 8.0, 1.2 mM EDTA, 167 mM NaCl, 1.1% Triton X-100, 0.01% SDS). They were sonicated for 5 cycles 30s ON/OFF in a Bioruptor sonicator (Diagenode). Samples were centrifuged; clear supernatant is transferred to a 15 mL tube. Aliquot of supernatant was kept for input DNA extraction. Supernatant diluted with IP dilution buffer. Antibody was added to supernatant and incubated overnight on a rocker at 4°C. Pre-washed DynaBeads were added to the mix and incubated for 1 hour at 4°C on a rocker. Beads were washed with low salt buffer (0.1% SDS, 1% Triton X-100, 2 mM EDTA, 20mM Tris pH 8.0, 150 mM NaCl), high salt buffer (0.1% SDS, 1% Triton X-100, 2 mM EDTA, 20 mM Tris pH 8.0, 500 mM NaCl) and 1X TE buffer (10 mM Tris pH 8.0, 1 mM EDTA). Beads are then incubated with elution buffer (0.1 M NaHCO_3_, 1% SDS) at room temperature for 15 minutes and eluted chromatin was reverse cross-linked by adding de-crosslinking buffer (2 M NaCl, 0.1 M EDTA, 0.4M Tris pH 7.5) and incubated at 65°C overnight on a rotator. Proteinase K (50 mg/mL) was used to digest proteins, for 2 hours at 50°C on a rocker. DNA was isolated with a Monarch PCR and DNA Cleanup kit. Sequencing and library prep information was previously described ^20^.

### ChIP-seq analysis

UPF1 and Ser2 RNA Polymerase II ChIP-seq datasets (GEO: GSE116806) were initially analysed as previously described ^20^. Additional analysis was done with Polymerase II ChIP-seq public datasets ^61^(GEO:GSE18643). Files were trimmed using TRIMMOMATIC (v0.39)^62^ to remove low quality reads, and adapter sequences. Output files were aligned to the genome using Bowtie2 v2.5.1 after the construction of a Bowtie2 index of *D. melanogaster* genome BDGP6.46 (Ensembl release 111). The aligned SAM files were converted to BAM files which were sorted and indexed using SAMtools v1.19.2. BAM files were converted to BEDgraph files for visualisation in the genome browser IGV ^63^ using the command genomeCoverageBed command and the options -bga and -ibam from the deepTools v3.5.2 suite ^64^.

### Genomic DNA extraction and quantification

In all instances of genomic DNA (gDNA) extraction, the GeneJET Genomic DNA Purification Kit (K0721, ThermoFisher Scientific) was used. For tissue samples, 20 samples were dissected in triplicate in 1X PBS, centrifuged briefly and PBS eluted off. S2 cells (2 x 10^7^) were centrifuged to form a pellet and washed twice, briefly with 1X PBS. Following the centrifugation and removal of the supernatant from samples, extraction was performed using the protocol recommended for: Mammalian Tissue and Rodent Tail Genomic DNA Purification Protocol. Tissue samples were homogenised with pestles after pelleting to the bottom of the tube by centrifugation, whilst S2 cells were resuspended in the kit-provided digestion solution, directly. A minimum of two hours was allowed for digestion of the sample; testes samples required a longer incubation of 5 hours for complete dissolution. At this point DNA was purified as described in the kit protocol.

To assess mtDNA copy number, extracted gDNA was diluted 1:50 in nuclease-free dH_2_O, prior to addition to the qPCR reaction. Each reaction consisted of 3 µL of a primer mix (final concentration of 400 nM), 2 µL of diluted gDNA and 5 µL qPCRBIO SyGreen Blue Mix Hi-ROX (PB20.16-50, PCR Biosystems Ltd).

### RNA extraction and quantification

Testes were dissected in 1X PBS, and transferred into ice-cold 1X PBS, the tissue was centrifuged at 1000 g for 30 seconds, the supernatant eluted off and directly resuspended in 1 mL TRIzol^TM^ Reagent (15596029, ThermoFisher Scientific). Tissue was incubated for 15 minutes at room temperature, vortexed for 1 minute, and 200 µL chloroform was added. Tubes were inverted by hand for 15 seconds until there was a homogenous suspension. Samples were centrifuged for 20 minutes at 15 000 g at 4°C. The top aqueous phase was eluted and collected in a sterile microcentrifuge tube. 1 µL RNase-free molecular grade glycogen (10901393001, Merck), and an equal amount ice-cold isopropanol was added. Samples were incubated on ice for 20 minutes to allow precipitation of RNA and centrifuged for 20 minutes at 15 000 g. The supernatant was discarded, and the pellet was washed twice in 75% ethanol, with 5-minute centrifuge spins. The pellet is allowed to dry and is resuspended in 20 µL DEPC-treated water.

Quality and quantification of RNA following dissolution was assessed and concentration measured using a spectrophotometer (Nanodrop-8000, ND-8000-GL, ThermoFisher Scientific). Quantification of RNA targets was assessed by RT-qPCR. Synthesis of cDNA used the qScript cDNA synthesis kit (7331174, QuantaBio), 200 ng of RNA was added to the 1X master mix alongside 1X qScript reverse transcriptase and the final volume was adjusted to 20 µL with the addition of nuclease-free water. Samples were placed in a thermocycler for the following steps: 22°C for 5 minutes, 42°C for 1 hour, and 82°C for 5 minutes. The product of cDNA synthesis was diluted 1:25 in nuclease-free water and proceeded with RT-qPCR quantification. The same reaction was prepared as for gDNA quantification, mentioned prior (primer concentration was 400 nM).

### RNA-sequencing and analysis

Prior to sequencing, PolyA mRNA was enriched. RNA libraries were then prepared commercially by Novogene and commercially assessed for quality. Sequencing of RNA was paired end of 150 bp in length and sequenced on Illumina NovaSeq platforms.

FASTQ files were trimmed using TRIMMOMATIC PE (v0.39) ^62^. Once trimmed, files were aligned to the *D. melanogaster* genome (BDGP6.56, FlyBase release FB2024_01), using HISAT2 (v2.2.1), after the construction of an HISAT2 index with the build function ^65^. SAMtools (v1.19.2) was used to convert the SAM files and to sort the BAM files based on genomic coordinates ^66^. Read counts were extracted using HTSeq count function (v0.13.5)^67^, and differential expression analysis was conducted using DESeq2 in R ^68^. To examine gene sets, values were extracted and plotted with gene lists curated on FlyBase for spermatogenesis (GO:0009283) and mitochondrion (GO:0005739). The testes RNA-seq datasets corresponding to two biological replicates of both UPF1^RNAi(1)^ and UPF1^RNAi(2)^ were processed separately, but the four counts files were averaged in the DESeq2 analysis, against the wild type replicates.

Gene Ontology (GO) analysis was performed using geneontology.org, downloaded results were plotted using R. Background gene lists were defined as all genes detected by sequencing, irrespective of log fold change.

### Primer list

Primers for dsRNA generation:

UPF1: F 5′-CCAGCATCACCAACGATCTG-3′, R 5′-AGTTGTTGCACATGACCACC-3

UPF2: F 5′-AGTTCCCGATTCGCATACCT-3′, R 5′-TCGCTTGCTCTCGATCTTCT-3′

UPF3: F 5′-CCTCCAAGGACATAGGCGAGG-3′, R 5′-CTCTCGATGGTGTTCACCTTGC-3′

qPCR primers:

UPF2: F 5′-TCAAGGCCACCGAGAAGATT-3′, R 5′-AATGCACCAGAACAACACCC-3′

UPF3: F 5′-CTCCCTTCCAGTGCTTCCTT-3′, R 5′-GGAGGGGTGTAGACTTGACC-3′

RpL32: F: 5′-CTCCCTTCCAGTGCTTCCTT-3′, R: 5′-GGAGGGGTGTAGACTTGACC-3′

Cox1: F 5′-TCCTGATATAGCATTCCCACGA-3′, R 5′-CCACCATGAGCAATTCCAGC-3′

ND4: F 5′-GCTCATGTTGAAGCTCCAGT-3′, R 5′-AAGCCTTTAAATCAGTTTGACGT-3′

Cyt-b: F 5′-CACCTGCCCATATTCAACCAGAATG-3′, R 5′-CAACTGGTCGAGCTCCAATTCAAG-3′

16S lrRNA: F 5′-AGTCTAACCTGCCCACTGAA-3′, R 5′-TCCAACCATTCATTCCAGCC-3′

### High Resolution Respirometry

High resolution respirometry was assessed in homogenised salivary glands from wild type and *Fkh>Upf1^RNAi(2)^* using an Oxygraph-2k (Oroboros Instruments, Innsbruck, Austria). Salivary glands from wild type and UPF1KD were added to each Oxygraph chamber containing 2 ml mitochondrial respiration buffer (MiR05,) at 25°C. Instrumental background oxygen flux was corrected for each sensor, accounting for sensor oxygen consumption and oxygen diffusion between the medium and chamber boundaries. Oxygen concentration in the chambers was maintained between 50-250 μM to avoid oxygen limitation of gland respiration. To establish mitochondrial electron transfer complex functions a substrate-uncoupler-inhibitor titration (SUIT) protocol was run on each sample. In brief, 2 mM malate plus 5 mM pyruvate were added sequentially to support NAD-linked substrate electron transport at Complex I (CI). Any oxygen consumption in the absence of ADP and ATP, was regarded as proton leak (LEAK respiration, L). ATP-coupled respiration was stimulated by 2.5 mM ADP, followed by 10 μM cytochrome c to test the integrity of the outer mitochondrial membrane. Further addition of the 10 mM glutamate was added to yield reduced nicotinamide adenine dinucleotide (NADH), which feeds electrons into complex I (NADH-ubiquinone oxidoreductase). Next, 10 mM Succinate was added to stimulate complex II respiration. To examine electron transfer capacity titrations of the uncoupler carbonylcyanide p-trifluoromethoxyphenyl- hydrazone (FCCP) was added sequentially at 0.5 μM intervals. Non-mitochondrial respiration was established by the sequential addition of 1 μM rotenone. Finally, complex IV capacity was assessed by adding 2 mM ascorbate and 0.5 mM TMPD - TMPD is a complex IV-specific electron donor, while ascorbate ensures that TMPD is reduced and continues to donate electrons to build a linear rate of complex IV activity. Specific autooxidation of ascorbate and TMPD was corrected for by inhibiting complex IV with azide (200 mM).

## Data availability

All RNA-seq datasets generated, and corresponding gene expression raw count data can be found on the gene expression omnibus (GEO) repository, accession number GSE266138.

## Acknowledgments

We would like to thank Hansong Ma and Ason Chiang for insightful discussions. We would like to thank the following for fly lines provided for this study. Helen White-Cooper (University of Cardiff) for sending us *bamGAL4:VP16*, Alicia Hidalgo (University of Birmingham) for providing us the *MhcGAL4* line for adult muscle experiments, and Hansong Ma (University of Birmingham) for providing, *Dj-GFP*, and the unpublished *ubi-mito-mCherry* line. The authors are also grateful to the staff of the Birmingham Advanced Light Microscopy (BALM) facilities, at the University of Birmingham. Also, the STED super-resolution microscopy of SATHI-Central Discovery Centre, and Confocal microscopy and advanced light microscopy (Interdisciplinary school of Life Sciences, Institute of Sciences, Banaras Hindu University, India) facilities. We would also like to thank the Biological Sciences Research Council [BBSRC BB/S017984/1, The Midlands Integrative Biosciences (Doctoral) Training Partnership (MIBTP) studentship to HLD], and Science and Engineering Research Board, CRG/2022/002767; Department of Biotechnology, Ministry of Science and Technology India, BT/RHD/35/02/2006, for funding this research.

---

**Figure S1.**
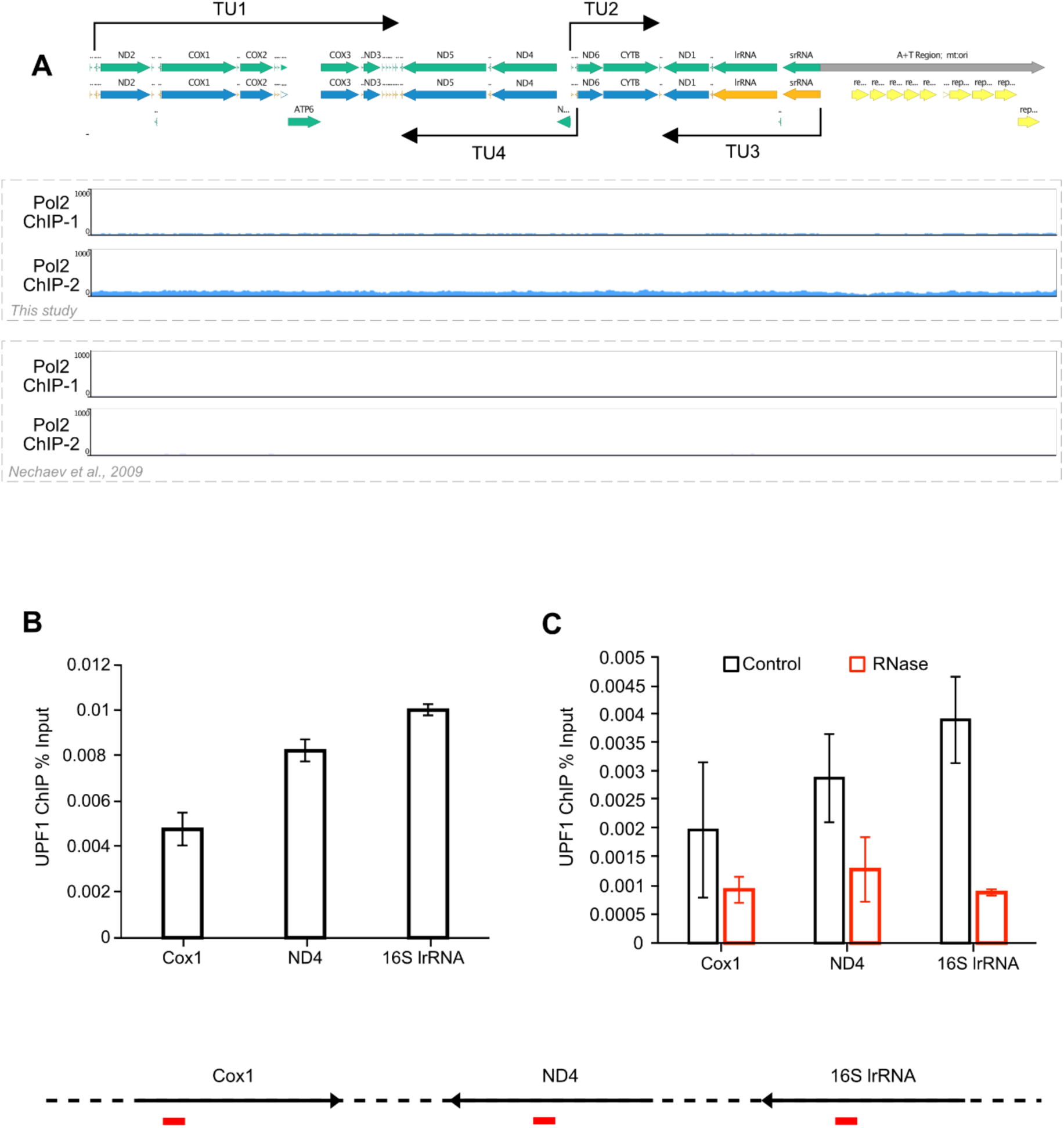
UPF1 ChIP-qPCR validation and RNase sensitivity evidence. (**A**) Top panel shows the annotated linear map of the mitochondrial genome. First two panels below show lack of mtDNA enrichment in two independent Pol II ChIP-Seq experiments ^25^. The next two rows show lack of enrichment in two Pol II ChIP-Seq datasets from a different laboratory ^61^. (**B**) ChIP-qPCR examination of UPF1 association with the three mtDNA gene regions indicated, Cox1, ND4 and lrRNA. The locations of the three primer pairs are indicated in red in the schematic below. (**C**) UPF1 ChIP-qPCR of the three mtDNA genes with, or without RNase treatment of the chromatin prior the IP (Material and Methods), error bars represent the standard error of the mean (SEM). The values are based on two separate ChIP experiments.

**Figure S2.**
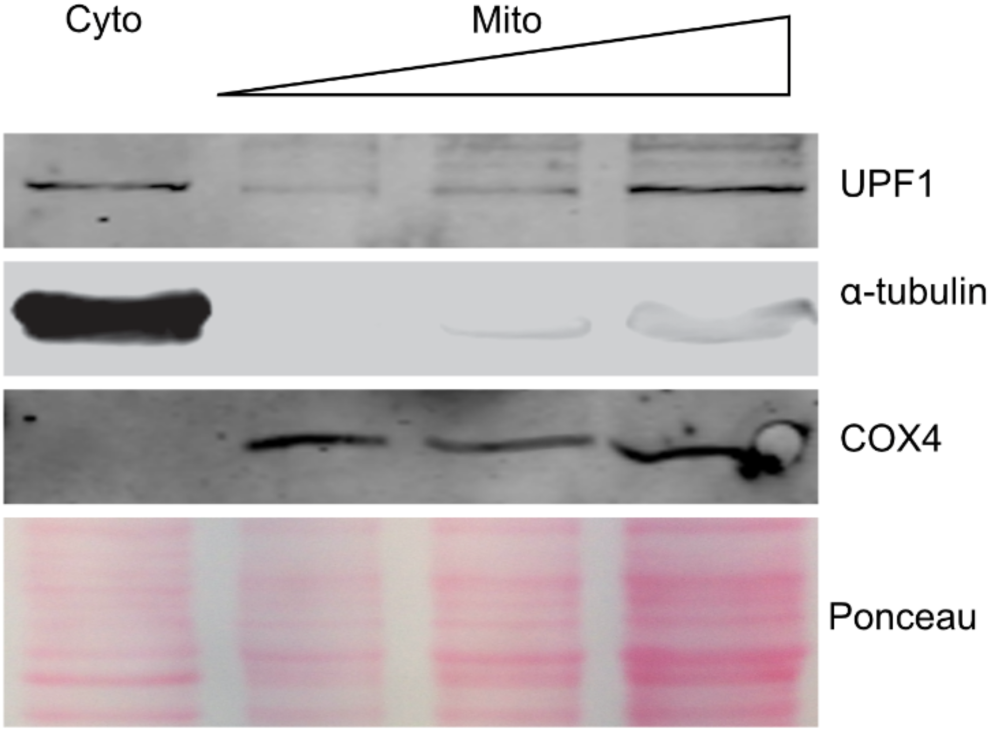
UPF1 copurifies with COX4 in S2 cells. Co-fractionation western blot experiment of cytoplasmic (Cyto) and mitochondrial (Mito) fractions. Both fractions were stained for UPF1 alongside α-tubulin and COX4 as controls for the cytoplasm and mitochondria, respectively. Ponceau staining of the membrane serves as a loading control.

**Figure S3.**
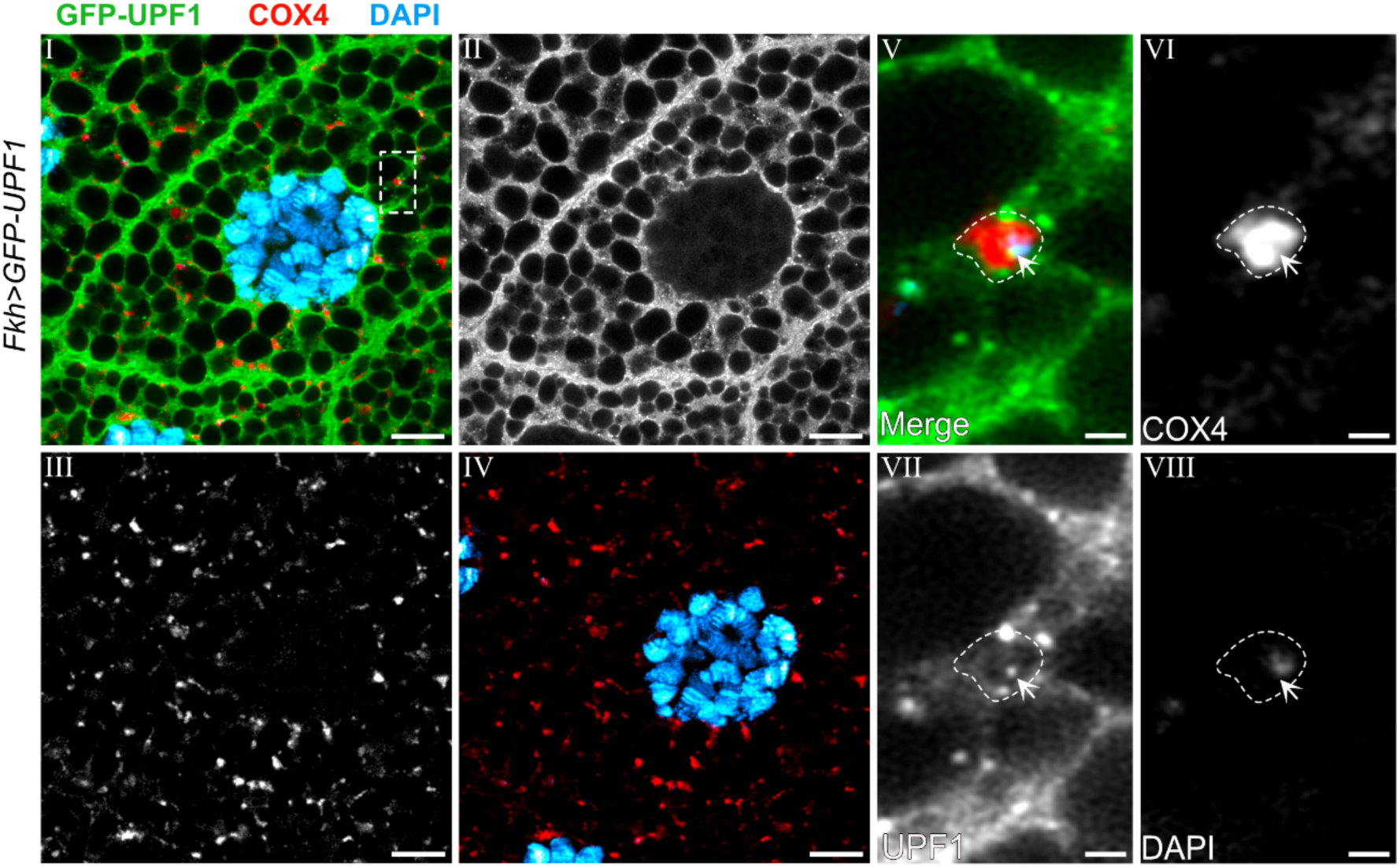
A portion of GFP-tagged UPF1 localises to mitochondria in salivary glands. Fluorescence immunolocalisation of *FkhGAL4* driven expression of GFP-UPF1 (green) and immunostaining of endogenous COX4 (red). DAPI (blue) represents nuclear and mtDNA in third instar salivary glands. Panel I shows a merged image of all three channels, UPF1 (panel II), COX4 (panel III). Panel IV displays both COX4 and DAPI. The boxed region shown in panel I is magnified in panels V-VIII, as a merged image (V) and grey for COX4 (VI), UPF1 (VII) and DAPI (VIII). Arrows point to foci of UPF1 adjacent to DAPI-stained regions of the mitochondria. Scale bar represents 10 µm.

**Figure S4.**
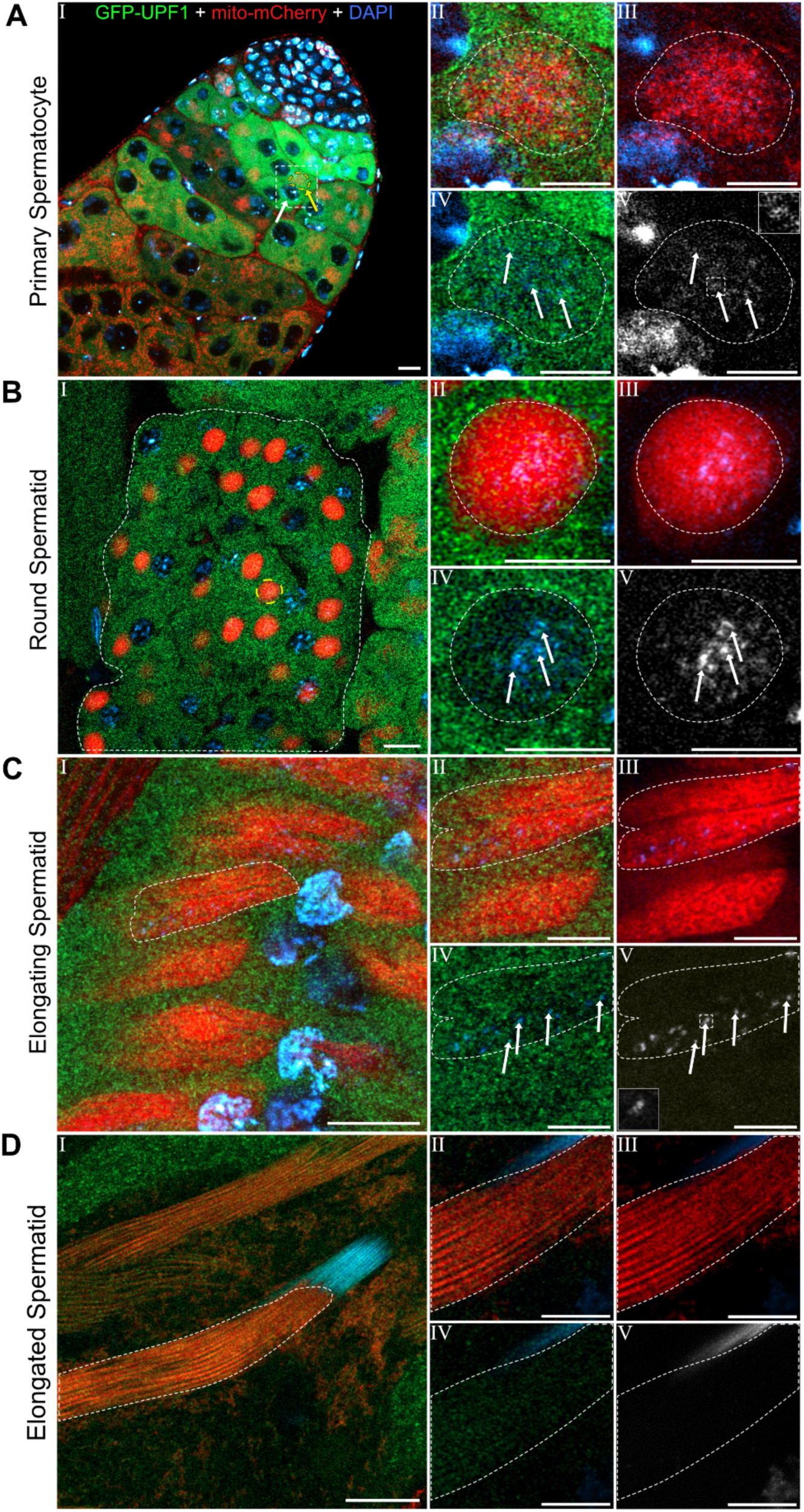
GFP-UPF1 also localises to mitochondria in developing spermatids. Fluorescent localization of GFP-UPF1 (green), ubiquitously expressed mCherry-mito (red) as the mitochondrial marker, and DAPI (blue) to stain nuclear and mitochondria DNA. (**A**) Primary spermatocytes. Panel I shows a merged image of the three channels, with a primary spermatocyte outlined by the white dotted box, the white arrow corresponds to the nucleus, and the yellow arrow marks the Mitoball. Panel II represents the merged magnified view of the outlined primary spermatocyte in panel I, the mitoball is outlined by the dashed white line. Panel III – IV represent mCherry-mito (panel III) and GFP-UPF1 (panel IV) together with DAPI. Panel V shows DAPI alone, with a magnified inset panel showing an example of a nucleoid region (top right). Arrows in panels IV and V indicate strong UPF1 signals proximal to DAPI signals. (**B**) Shows a cyst with round-stage spermatids. Magnified views of the nebenkern outlined in yellow in panel I are shown in panels II-V. White arrows in panels IV and V indicate GFP-UPF1 dots adjacent to DAPI in the centre nebenkern. (**C**) GFP-UPF1 localisation in elongating spermatids. Magnified view of a portion of the area (major and minor mitochondrial derivatives) outlined by the dashed line in panel I are shown in panels II – V. Arrows in panels IV and V show strong signals of GFP-UPF1 adjacent to DAPI, magnification inset of DAPI is seen in panel V (bottom left). Panel V shows DAPI alone. Arrows in panels IV and V indicate strong GFP-UPF1 signals proximal to DAPI signal. (**D**) Imaging of elongated spermatids, which show absence of GFP-UPF1. Panel I shows the outlined region which is magnified in panels II-V which show absence of GFP-UPF1 and DAPI foci in the spermatid tail, panels IV-V. Scale bars in A-D, panel I represent 10 µm and panels II-IV represent 5 µm.

**Figure S5.**
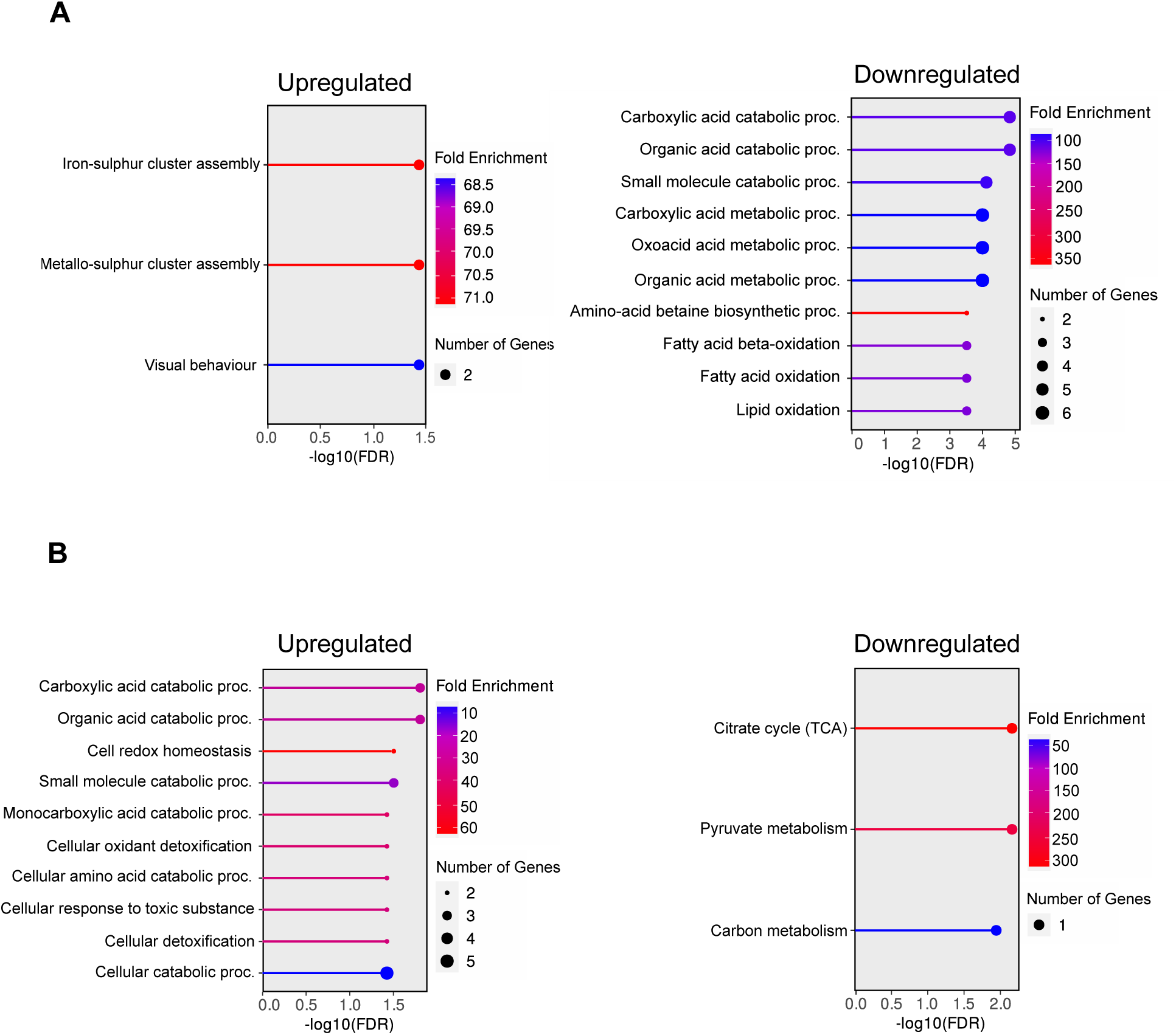
Mitochondrial-associated biological processes are not disrupted in UPF2 and UPF3KD salivary glands. Gene ontology analysis of UPF2KD (**A**) and UPF3KD (**B**) salivary glands, the left and right panels represent the upregulated and downregulated biological processes for each genotype, respectively. The node size corresponds to the number of genes associated with the GO term, colour corresponds to the fold enrichment, from blue (lowest) to red (highest), only pathways with a -log10 enrichment FDR <0.05 are shown. Pathways are ordered by -log10(FDR). Upregulated GO terms are as follows: UPF2KD: GO:0016226, GO:0031163, GO:0007632; UPF3KD: GO:0046395, GO:0016054, GO:0045454, GO:0044282, GO:0072329, GO:0098869, GO:0009063, GO:0097237, GO:1990748, GO:0009056. Downregulated GO terms are as follows: UPF2KD: GO:0046395, GO:0016054, GO:0044282, GO:0019752, GO:0043436, GO:0006082, GO:0006578, GO:0006635, GO:0019395, GO:0034440; UPF3KD: GO:0006099, GO:0006090, GO:0015976. GO terms are listed in order of appearance (top to bottom).

## References

1. Bhattacharya, A., Czaplinski, K., Trifillis, P., He, F., Jacobson, A., and Peltz, S.W. (2000). Characterization of the biochemical properties of the human Upf1 gene product that is involved in nonsense-mediated mRNA decay. RNA 6, 1226–1235.

2. Chakrabarti, S., Jayachandran, U., Bonneau, F., Fiorini, F., Basquin, C., Domcke, S., Le Hir, H., and Conti, E. (2011). Molecular mechanisms for the RNA-dependent ATPase activity of Upf1 and its regulation by Upf2. Mol Cell 41, 693–703. 10.1016/j.molcel.2011.02.010.

3. Czaplinski, K., Weng, Y., Hagan, K.W., and Peltz, S.W. (1995). Purification and characterization of the Upf1 protein - a factor involved in translation and messenger RNA degradation. RNA 1, 610–623.

4. Fiorini, F., Bagchi, D., Le Hir, H., and Croquette, V. (2015). Human Upf1 is a highly processive RNA helicase and translocase with RNP remodelling activities. Nature communications 6, 7581. 10.1038/ncomms8581.

5. Hogg, J.R., and Goff, S.P. (2010). Upf1 senses 3’UTR length to potentiate mRNA decay. Cell 143, 379–389. 10.1016/j.cell.2010.10.005.

6. Hurt, J.A., Robertson, A.D., and Burge, C.B. (2013). Global analyses of UPF1 binding and function reveal expanded scope of nonsense-mediated mRNA decay. Genome Res 23, 1636–1650. 10.1101/gr.157354.113.

7. Zund, D., Gruber, A.R., Zavolan, M., and Muhlemann, O. (2013). Translation-dependent displacement of UPF1 from coding sequences causes its enrichment in 3’ UTRs. Nat Struct Mol Biol 20, 936–943. 10.1038/nsmb.2635.

8. Brogna, S., and Wen, J. (2009). Nonsense-mediated mRNA decay (NMD) mechanisms. Nat Struct Mol Biol 16, 107–113. 10.1038/nsmb.1550.

9. Fatscher, T., Boehm, V., and Gehring, N.H. (2015). Mechanism, factors, and physiological role of nonsense-mediated mRNA decay. Cell Mol Life Sci 72, 4523–4544. 10.1007/s00018-015-2017-9.

10. Goetz, A.E., and Wilkinson, M. (2017). Stress and the nonsense-mediated RNA decay pathway. Cell Mol Life Sci 74, 3509–3531. 10.1007/s00018-017-2537-6.

11. He, F., and Jacobson, A. (2015). Nonsense-Mediated mRNA Decay: Degradation of Defective Transcripts Is Only Part of the Story. Annu Rev Genet. 10.1146/annurev-genet-112414-054639.

12. Hug, N., Longman, D., and Caceres, J.F. (2016). Mechanism and regulation of the nonsense-mediated decay pathway. Nucleic Acids Res 44, 1483–1495. 10.1093/nar/gkw010.

13. Karousis, E.D., Nasif, S., and Muhlemann, O. (2016). Nonsense-mediated mRNA decay: novel mechanistic insights and biological impact. Wiley Interdiscip Rev RNA 7, 661–682. 10.1002/wrna.1357.

14. Kim, Y.K., and Maquat, L.E. (2019). UPFront and center in RNA decay: UPF1 in nonsense-mediated mRNA decay and beyond. RNA 25, 407–422. 10.1261/rna.070136.118.

15. Lykke-Andersen, S., and Jensen, T.H. (2015). Nonsense-mediated mRNA decay: an intricate machinery that shapes transcriptomes. Nat Rev Mol Cell Biol 16, 665–677. 10.1038/nrm4063.

16. Brogna, S., McLeod, T., and Petric, M. (2016). The Meaning of NMD: Translate or Perish. Trends Genet 32, 395–407. 10.1016/j.tig.2016.04.007.

17. Ajamian, L., Abel, K., Rao, S., Vyboh, K., Garcia-de-Gracia, F., Soto-Rifo, R., Kulozik, A.E., Gehring, N.H., and Mouland, A.J. (2015). HIV-1 Recruits UPF1 but Excludes UPF2 to Promote Nucleocytoplasmic Export of the Genomic RNA. Biomolecules 5, 2808–2839. 10.3390/biom5042808.

18. Mendell, J.T., ap Rhys, C.M., and Dietz, H.C. (2002). Separable roles for rent1/hUpf1 in altered splicing and decay of nonsense transcripts. Science 298, 419–422.

19. Nasif, S., Eberle, A.B., Schranz, K., Hadorn, R., Chakrabarti, S., and Muhlemann, O. (2025). UPF1 shuttles between nucleus and cytoplasm independently of its RNA-binding and ATPase activities. RNA 31, 1872–1885. 10.1261/rna.080476.125.

20. Singh, A.K., Choudhury, S.R., De, S., Zhang, J., Kissane, S., Dwivedi, V., Ramanathan, P., Orsini, L., Hebenstreit, D., and Brogna, S. (2019). The RNA helicase UPF1 associates with mRNAs co-transcriptionally and is required for the release of mRNAs from transcription sites. Elife 2019;*8*:e41444. 10.7554/eLife.41444.

21. De, S., Varsally, W., Falciani, F., and Brogna, S. (2011). Ribosomal proteins’ association with transcription sites peaks at tRNA genes in Schizosaccharomyces pombe. RNA 17, 1713–1726. 10.1261/rna.2808411.

22. Azzalin, C.M., and Lingner, J. (2006). The human RNA surveillance factor UPF1 is required for S phase progression and genome stability. Curr Biol 16, 433–439.

23. Azzalin, C.M., Reichenbach, P., Khoriauli, L., Giulotto, E., and Lingner, J. (2007). Telomeric repeat containing RNA and RNA surveillance factors at mammalian chromosome ends. Science 318, 798–801.

24. De, S., Edwards, D.M., Dwivedi, V., Wang, J., Varsally, W., Dixon, H.L., Singh, A.K., Owuamalam, P.O., Wright, M.T., Summers, R.P., et al. (2022). Genome-wide chromosomal association of Upf1 is linked to Pol II transcription in Schizosaccharomyces pombe. Nucleic Acids Res 50, 350–367. 10.1093/nar/gkab1249.

25. Singh, A.K., Zhang, J., Hebenstreit, D., and Brogna, S. (2020). Evidence of slightly increased Pol II pausing in UPF1-depleted Drosophila melanogaster cells. MicroPubl Biol 2020. 10.17912/micropub.biology.000319.

26. Zaepfel, B.L., Zhang, Z., Maulding, K., Coyne, A.N., Cheng, W., Hayes, L.R., Lloyd, T.E., Sun, S., and Rothstein, J.D. (2021). UPF1 reduces C9orf72 HRE-induced neurotoxicity in the absence of nonsense-mediated decay dysfunction. Cell Rep 34, 108925. 10.1016/j.celrep.2021.108925.

27. Zuniga, G., Levy, S., Ramirez, P., De Mange, J., Gonzalez, E., Gamez, M., and Frost, B. (2023). Tau-induced deficits in nonsense-mediated mRNA decay contribute to neurodegeneration. Alzheimers Dement 19, 405–420. 10.1002/alz.12653.

28. Chen, H., Ma, J., Kong, F., Song, N., Wang, C., and Ma, X. (2022). UPF1 contributes to the maintenance of endometrial cancer stem cell phenotype by stabilizing LINC00963. Cell Death Dis 13, 257. 10.1038/s41419-022-04707-x.

29. Staszewski, J., Lazarewicz, N., Konczak, J., Migdal, I., and Maciaszczyk-Dziubinska, E. (2023). UPF1-From mRNA Degradation to Human Disorders. Cells 12. 10.3390/cells12030419.

30. Petric Howe, M., and Patani, R. (2023). Nonsense-mediated mRNA decay in neuronal physiology and neurodegeneration. Trends Neurosci 46, 879–892. 10.1016/j.tins.2023.07.001.

31. Su, W., Kochen Rossi, J., Nuevo-Tapioles, C., Chen, T., Kawaler, E., Branco, C., Wong, K.-k., Simeone, D.M., Gardner, L.B., and Philips, M.R. (2024). UPF1 deficiency enhances mitochondrial ROS which promotes an immunosuppressive microenvironment in pancreatic ductal adenocarcinoma. Proceedings of the National Academy of Sciences 121, e2401996121.

32. Culbertson, M.R. (1999). RNA surveillance - unforeseen consequences for gene expression, inherited genetic disorders and cancer. Trends in Genetics 15, 74–80.

33. Leeds, P., Peltz, S.W., Jacobson, A., and Culbertson, M.R. (1991). The product of the yeast Upf1 gene is required for rapid turnover of messenger RNAs containing a premature translational termination codon. Genes & development 5, 2303–2314.

34. Altamura, N., Groudinsky, O., Dujardin, G., and Slonimski, P.P. (1992). Nam7 nuclear gene encodes a novel member of a family of helicases with a Zn-ligand motif and is involved in mitochondrial functions in *Saccharomyces cerevisiae*. Journal of Molecular Biology 224, 575–587.

35. Asher, E.B., Groudinsky, O., Dujardin, G., Altamura, N., Kermorgant, M., and Slonimski, P.P. (1989). Novel class of nuclear genes involved in both mRNA splicing and protein synthesis in Saccharomyces cerevisiae mitochondria. Mol Gen Genet 215, 517–528. 10.1007/BF00427051.

36. de Pinto, B., Lippolis, R., Castaldo, R., and Altamura, N. (2004). Overexpression of Upf1p compensates for mitochondrial splicing deficiency independently of its role in mRNA surveillance. Mol Microbiol 51, 1129–1142. 10.1046/j.1365-2958.2003.03889.x.

37. Lewis, D.L., Farr, C.L., Farquhar, A.L., and Kaguni, L.S. (1994). Sequence, organization, and evolution of the A+T region of Drosophila melanogaster mitochondrial DNA. Mol Biol Evol 11, 523–538. 10.1093/oxfordjournals.molbev.a040132.

38. Holt, I.J., and Reyes, A. (2012). Human mitochondrial DNA replication. Cold Spring Harb Perspect Biol 4. 10.1101/cshperspect.a012971.

39. Joers, P., and Jacobs, H.T. (2013). Analysis of replication intermediates indicates that Drosophila melanogaster mitochondrial DNA replicates by a strand-coupled theta mechanism. PLoS One 8, e53249. 10.1371/journal.pone.0053249.

40. Torres, T.T., Dolezal, M., Schlotterer, C., and Ottenwalder, B. (2009). Expression profiling of Drosophila mitochondrial genes via deep mRNA sequencing. Nucleic Acids Res 37, 7509–7518. 10.1093/nar/gkp856.

41. Bogenhagen, D.F. (2012). Mitochondrial DNA nucleoid structure. Bba-Gene Regul Mech 1819, 914–920. 10.1016/j.bbagrm.2011.11.005.

42. Li, A.Y.Z., Di, Y., Rathore, S., Chiang, A.C., Jezek, J., and Ma, H. (2023). Milton assembles large mitochondrial clusters, mitoballs, to sustain spermatogenesis. Proc Natl Acad Sci U S A 120, e2306073120. 10.1073/pnas.2306073120.

43. Hales, K.G., and Fuller, M.T. (1997). Developmentally regulated mitochondrial fusion mediated by a conserved, novel, predicted GTPase. Cell 90, 121–129. 10.1016/s0092-8674(00)80319-0.

44. Lee, H.S., Simon, J.A., and Lis, J.T. (1988). Structure and expression of ubiquitin genes of Drosophila melanogaster. Mol Cell Biol 8, 4727–4735. 10.1128/mcb.8.11.4727-4735.1988.

45. Patel, M.R. (2017). Inheritance: Male mtDNA Just Can’t Catch a Break. Curr Biol 27, R264–R266. 10.1016/j.cub.2017.02.057.

46. White-Cooper, H. (2012). Tissue, cell type and stage-specific ectopic gene expression and RNAi induction in the Drosophila testis. Spermatogenesis 2, 11–22. 10.4161/spmg.19088.

47. Bezawork-Geleta, A., Rohlena, J., Dong, L., Pacak, K., and Neuzil, J. (2017). Mitochondrial complex II: at the crossroads. Trends in biochemical sciences 42, 312–325.

48. Zhou, Z., Ma, A., Moore, T.M., Wolf, D.M., Yang, N., Tran, P., Segawa, M., Strumwasser, A.R., Ren, W., and Fu, K. (2024). Drp1 controls complex II assembly and skeletal muscle metabolism by Sdhaf2 action on mitochondria. Science advances 10, eadl0389.

49. Rorbach, J., Richter, R., Wessels, H.J., Wydro, M., Pekalski, M., Farhoud, M., Kuhl, I., Gaisne, M., Bonnefoy, N., Smeitink, J.A., et al. (2008). The human mitochondrial ribosome recycling factor is essential for cell viability. Nucleic Acids Res 36, 5787–5799. 10.1093/nar/gkn576.

50. Gray, L.R., Tompkins, S.C., and Taylor, E.B. (2014). Regulation of pyruvate metabolism and human disease. Cell Mol Life Sci 71, 2577–2604. 10.1007/s00018-013-1539-2.

51. Bhuiyan, S.H., Bordet, G., Bamgbose, G., and Tulin, A.V. (2023). The Drosophila gene encoding JIG protein (CG14850) is critical for CrebA nuclear trafficking during development. Nucleic Acids Res 51, 5647–5660. 10.1093/nar/gkad343.

52. Chacinska, A., Koehler, C.M., Milenkovic, D., Lithgow, T., and Pfanner, N. (2009). Importing mitochondrial proteins: machineries and mechanisms. Cell 138, 628–644. 10.1016/j.cell.2009.08.005.

53. Fukasawa, Y., Tsuji, J., Fu, S.C., Tomii, K., Horton, P., and Imai, K. (2015). MitoFates: improved prediction of mitochondrial targeting sequences and their cleavage sites. Mol Cell Proteomics 14, 1113–1126. 10.1074/mcp.M114.043083.

54. Savojardo, C., Bruciaferri, N., Tartari, G., Martelli, P.L., and Casadio, R. (2020). DeepMito: accurate prediction of protein sub-mitochondrial localization using convolutional neural networks. Bioinformatics 36, 56–64. 10.1093/bioinformatics/btz512.

55. Bader, G., Enkler, L., Araiso, Y., Hemmerle, M., Binko, K., Baranowska, E., De Craene, J.O., Ruer-Laventie, J., Pieters, J., Tribouillard-Tanvier, D., et al. (2020). Assigning mitochondrial localization of dual localized proteins using a yeast Bi-Genomic Mitochondrial-Split-GFP. Elife 9. 10.7554/eLife.56649.

56. Henderson, K.D., and Andrew, D.J. (2000). Regulation and function of Scr, exd, and hth in the Drosophila salivary gland. Dev Biol 217, 362–374.

57. Schuster, C.M., Davis, G.W., Fetter, R.D., and Goodman, C.S. (1996). Genetic dissection of structural and functional components of synaptic plasticity. I. Fasciclin II controls synaptic stabilization and growth. Neuron 17, 641–654. 10.1016/s0896-6273(00)80197-x.

58. Singh, A.K., and Lakhotia, S.C. (2024). Visualizing dynamic organization of nuclei and mitochondria during spermatogenesis in Drosophila. In Experiments with Drosophila for Biology Courses, S.C. Lakhotia, ed. (Indian Academy of Sciences Bengaluru).

59. Tenenbaum, C.M., and Gavis, E.R. (2016). Removal of Drosophila Muscle Tissue from Larval Fillets for Immunofluorescence Analysis of Sensory Neurons and Epidermal Cells. J Vis Exp. 10.3791/54670.

60. Schindelin, J., Arganda-Carreras, I., Frise, E., Kaynig, V., Longair, M., Pietzsch, T., Preibisch, S., Rueden, C., Saalfeld, S., Schmid, B., et al. (2012). Fiji: an open-source platform for biological-image analysis. Nature methods 9, 676–682. 10.1038/nmeth.2019.

61. Nechaev, S., Fargo, D.C., dos Santos, G., Liu, L., Gao, Y., and Adelman, K. (2010). Global analysis of short RNAs reveals widespread promoter-proximal stalling and arrest of Pol II in Drosophila. Science 327, 335–338. 10.1126/science.1181421.

62. Bolger, A.M., Lohse, M., and Usadel, B. (2014). Trimmomatic: a flexible trimmer for Illumina sequence data. Bioinformatics 30, 2114–2120. 10.1093/bioinformatics/btu170.

63. Robinson, J.T., Thorvaldsdottir, H., Turner, D., and Mesirov, J.P. (2023). igv.js: an embeddable JavaScript implementation of the Integrative Genomics Viewer (IGV). Bioinformatics 39. 10.1093/bioinformatics/btac830.

64. Ramirez, F., Dundar, F., Diehl, S., Gruning, B.A., and Manke, T. (2014). deepTools: a flexible platform for exploring deep-sequencing data. Nucleic Acids Res 42, W187–191. 10.1093/nar/gku365.

65. Kim, D., Paggi, J.M., Park, C., Bennett, C., and Salzberg, S.L. (2019). Graph-based genome alignment and genotyping with HISAT2 and HISAT-genotype. Nature biotechnology 37, 907–915. 10.1038/s41587-019-0201-4.

66. Li, H., Handsaker, B., Wysoker, A., Fennell, T., Ruan, J., Homer, N., Marth, G., Abecasis, G., Durbin, R., and Genome Project Data Processing, S. (2009). The Sequence Alignment/Map format and SAMtools. Bioinformatics 25, 2078–2079. 10.1093/bioinformatics/btp352.

67. Anders, S., Pyl, P.T., and Huber, W. (2015). HTSeq--a Python framework to work with high-throughput sequencing data. Bioinformatics 31, 166–169. 10.1093/bioinformatics/btu638.

68. Love, M.I., Huber, W., and Anders, S. (2014). Moderated estimation of fold change and dispersion for RNA-seq data with DESeq2. Genome Biol 15, 550. 10.1186/s13059-014-0550-8.

